# Linking microenvironment modification to species interactions and demography in an alpine plant community

**DOI:** 10.1101/2022.06.21.497097

**Authors:** Courtenay A. Ray, Rozalia E. Kapas, Øystein H. Opedal, Benjamin W. Blonder

## Abstract

Individual plants can modify the microenvironment within their spatial neighborhood. However, the consequences of microenvironment modification for demography and species interactions remain unclear at the community scale. In a study of co-occurring alpine plants, we (1) determined the extent of species-specific microclimate modification by comparing temperature and soil moisture between vegetated and non-vegetated microsites for several focal species. We (2) determined how vital rates (survival, growth, fecundity) of all species varied in response to aboveground and belowground vegetative overlap with inter- and intraspecific neighbors as proxies for microenvironment modification. For (1), surface temperatures were buffered (lower maximums and higher minimums) and soil moisture was higher below the canopies of most species compared to non-vegetated areas. For (2), vegetative overlap predicted most vital rates, although the effect varied depending on whether aboveground or belowground overlap was considered. Vital rate response to microenvironment-modification proxies (vegetative overlap) was also frequently context dependent with respect to plant size and macroclimate. Microenvironment modification and spatial overlapping of individuals are key drivers of demography and species interactions in this alpine community.

## Introduction

Species interactions affect vital rates (*i*.*e*. survival, growth, fecundity) and may scale up to impact species distributions, co-occurrence patterns, and abundances (Diamond 1975; Callaway & Walker 1997; Wisz *et al*. 2012; Ulrich *et al*. 2016; Wright *et al*. 2017). The strength and direction of species interactions and their effects on vital rates are increasingly well understood, yet generalization has been challenged by the dependence of these interactions on the abiotic and biotic context (Soliveres *et al*. 2015; Matías *et al*. 2018). For example, the outcome of species interactions may vary across macroclimatic conditions (*i*.*e*. regional climate) (Wright *et al*. 2015) or shift in the presence of other species (Saavedra *et al*. 2017). Context dependency creates challenges in scaling up from pairwise to multispecies interactions due to species-specific responses to environmental context (Barabás *et al*. 2018). An improved understanding of species interactions at the community scale and their sensitivity to context is needed to better understand community assembly, and would contribute toward more accurate forecasts of community response to environmental change (Ehrlén & Morris 2015; Opedal *et al*. 2020).

Modification of abiotic conditions (*i*.*e*. microenvironment modification) is a widespread mechanism of species interactions (Bertness & Callaway 1994). However, few generalities exist concerning which factors may influence modification strength and when microenvironment modification may be important for species interactions. One reason why microenvironment modification is difficult to generalize is that alterations of the abiotic environment influence individual plant performance as well as the environmental context of interactions between other plants. In plant communities, for example, a plant can shift resource availability for another plant (Breshears *et al*. 1998), change the diversity and availability of suitable niches (Stark *et al*. 2017), and through both processes generate performance feedbacks on the modifying species (Pugnaire *et al*. 1996). The importance of microenvironment modification as a species interaction may also increase under harsher macroclimatic conditions such as those associated with decreased precipitation (Wright *et al*. 2015). Such variable effects of microenvironment modification represent a challenge for predicting when and how microenvironment modification as a species interaction shapes community assembly.

Scaling microenvironment modification effects from individuals to multispecies assemblages remains a challenge. Previous work addressing how microclimate modification affects vital rates often has used presence/absence comparisons, such as inside versus outside plants (*e*.*g*. Nuñez *et al*. 1999; Maestre *et al*. 2001; Nyakatya & McGeoch 2008). Such studies have shown that the effect of microenvironment modification on plant success often depends on the identity and proximity of the modifying species (Mack & Harper 1977; Rees *et al*. 1996). However, net interaction effects in multi-individual and multispecies assemblages are more complex and may depend on neighborhood biomass (Wright *et al*. 2015) and diversity (Wright *et al*. 2021), with many possible feedbacks. As spatial overlap among plants is common across ecosystems, an increased understanding of how plants interact when aggregated would provide insight into the drivers of plant distributions and occurrence patterns.

Aboveground and belowground microenvironment modification may influence species interactions. Aboveground, for example, plants may alter wind exposure, increase relative humidity, or alter ambient and surface temperatures (Breshears *et al*. 1998). In tundra environments, plants (particularly shrubs) may impact snow accumulation patterns (Sturm *et al*. 2005). Belowground, plant interactions via microenvironment modification may reflect alterations to resource availability, such as water (*e*.*g*. Zou *et al*. 2005), as well as effects of soil biota associations (Rodríguez-Echeverría *et al*. 2013). The strength and direction of species interactions may also differ above- and belowground. For example, plants may compete for nutrients belowground, while ameliorating microclimates aboveground (Klanderud 2005). While canopy size may predict the spatial area that a plant modifies aboveground, lateral root extent may predict modification belowground because it influences the distance over which a plant can acquire and redistribute resources. Due to differences in the relative size of above-versus belowground biomass, modification extent and the number and diversity of interacting partners may vary between these dimensions (Ottaviani *et al*. 2020). It is unclear the extent that above-or belowground overlap affect vital rates, as well as whether these effects are context dependent.

Alpine environments are an ideal setting to test how microenvironment modification affects plant performance as species interactions are known to be important for demography in this biome. For example, microenvironment modification is considered a common community assembly mechanism in the alpine as it may ameliorate abiotic conditions and increase the range of available niches (Holzapfel & Mahall 1999; Choler *et al*. 2001; Callaway *et al*. 2002; Drezner 2006). Other benefits of the alpine include low local species richness and extreme abiotic conditions where modification could have important demographic consequences (Körner 2003). By comparing vital rates in communities with high microenvironment modification and under different biotic contexts (*i*.*e*. varying species assemblages), we can gain insight into how microsite variation influences community assembly through effects on demography.

Here we use microenvironment and spatial distribution data from a long-term alpine plant community demography study. We determine how vital rates (survival, growth, fecundity) vary in response to vegetative overlap with other plants as a proxy for microenvironment modification, as well as how vital rates vary in response to environmental context, individual state, and utilization of above-or belowground spatial data. Vegetative overlap is a metric of co-occurrence and thus may be a good proxy of microenvironment modification through physical and physiological processes. We asked (Q1) whether and how much microenvironment modification occurs for several focal species in this community. We then asked (Q2) how vital rates respond to microenvironment modification, using vegetative overlap as a proxy of microenvironment modification, and whether the response depended on context (macroclimate) and an individual state parameter (focal plant size).

## Methods

### Data collection

#### Demographic census

We analyzed a demography dataset from an alpine community located in southwestern Colorado (38.978725°N, 107.042104°W, ∼3540 m above sea level). From 2014 to 2019, we tracked all individuals (> 2,300) of 18 species occurring on fifty 2 × 2 m permanent plots organized in a 5 × 10 array. The site is on a southeast-oriented ridgeline with a ∼20° slope. Soil development at this site is very limited, with only 5-10 cm of fine scree over bedrock. The growing season is short, as snow cover typically occurs from October to June. Vegetation is patchy with low cover (∼14.5% in 2014) (Figure 1). Eighteen perennial species are found at the site, including (in order of greatest abundance during the 2014 census) *Lupinus argenteus* (33.6%, Fabaceae), *Ivesia gordonii* (17.1%, Rosaceae), *Eriogonum umbellatum* (14.2%, Polygonaceae), *Elymus lanceolatus* (13.7%, Poaceae), *Heterotheca villosa* (7.7%, Asteraceae), and *Carex siccata* (3.4%, Cyperaceae). The site and species list are described in detail in Blonder *et al*. (2018).

**Figure 1:**
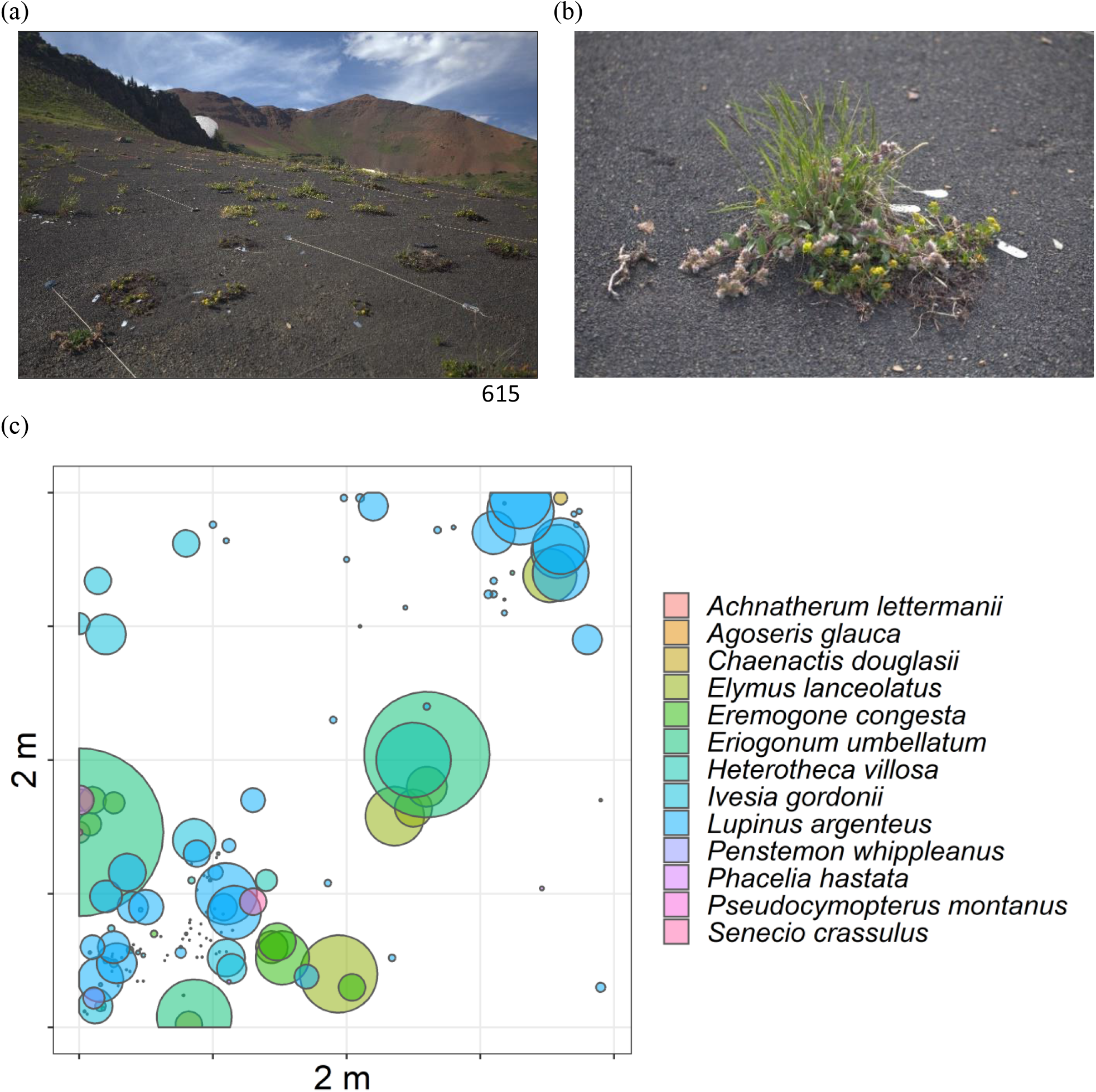
(a) A photograph of the study site during the growing season showing an open landscape with clustered vegetation. Plots are visible via white string boundaries; (b) An example vegetative cluster with high overlap featuring *E. umbellatum* (yellow flowers), *E. lanceolatus* (grass), *P. hastata* (pink flowers); (c) Circular polygons generated from census data from plot 20 in 2015. Polygons are scaled to the size of the plant with a distinct color for each taxon.

#### Microenvironment descriptors

To address Q1, we determined the existence and strength of microenvironment modification of surface temperature and soil moisture across several species. In 2016 and 2018, we measured surface temperatures (°C) across this site using iButton data loggers (Thermochron DS1921G, Maxim, San Jose, CA, USA). Data loggers were placed at soil level after being sealed in Parafilm (Bemis Company, Inc, Neenah, WI, USA) and grey duct tape for waterproofing and to roughly match the reflectance of the soil substrate, following the method of Stark *et al*. (2017). Temperatures measured by data loggers are not exactly equal to substrate temperature due to radiative loading, among other factors, especially in high light environments such as the alpine (Maclean *et al*. 2021). Deviations from true temperatures are expected to be greatest for temperature maximums which are typically during the afternoon versus temperature minimums which are typically during low light conditions, but measurements should be comparable across species. All temperature data were collected at 20-minute intervals. We summarized these data as maximum and minimum temperatures during deployment for each logger and each year.

We measured microenvironment within the canopy of a subset of species that were abundant in the plots and represented a range of growth forms, including erect dicotyledons (*Agoseris glauca* (Asteraceae), *L. argenteus, Senecio crassulus* (Asteraceae)), rosette dicotyledons (*Eremogone congesta* (Caryophyllaceae), *I. gordonii, Phacelia hastata* (Boraginaceae)), erect monocotyledons (*Achnatherum lettermanii* (Poaceae), *E. lanceolatus*), and deciduous dicotyledonous shrubs (*E. umbellatum, H. villosa*) (*sensu* (Webber *et al*. 1980; Callaghan *et al*. 2005)). We avoided measuring microenvironment within focal plants growing in clusters of multiple individuals. For plant-level measurements, data loggers were placed under the vegetative edge of each individual.

For 26 days during the growing season in 2016, we collected surface temperatures beneath 86 randomly selected focal plants of 9 common species (*A. lettermanii* (*n*=8), *A. glauca* (*n*=6), *E. lanceolatus* (*n*=13), *E. umbellatum* (*n*=11), *H. villosa* (*n*=10), *I. gordonii* (*n*=13), *L. argenteus* (*n*=10), *P. hastata* (*n*=8), *S. crassulus* (*n*=7)). Focal plants were selected randomly among censused individuals with vegetative diameters > 1 cm. Concurrently, we also collected temperature data at 91 plot corners, with data loggers at the upper left and bottom right corners of each plot. These plot corners were non-vegetated and selected to allow comparison with the plant-level measurements.

Using similar methods, in 2018, we augmented the data by measuring surface temperatures for 26 days under the canopy edge of 81 randomly selected individuals of 7 species: *E. congesta* (n=10), *E. lanceolatus* (*n*=12), *E. umbellatum* (*n*=14), *H. villosa* (*n*=10), *I. gordonii* (*n*=12), *L. argenteus* (*n*=13), *S. crassulus* (*n*=10). Six of these taxa were also sampled in 2016. However, in contrast to 2016, as a non-vegetated comparison, the iButtons were placed in a non-vegetated location 10 cm from the plant edge in a randomly selected direction. We used this paired vegetated versus non-vegetated sampling design in 2018 to reduce any effects of spatial distance on microsite microclimate.

We measured soil volumetric water content (%) at 3.8 cm depth on 9 August 2016 with a FieldScout TDR 100 probe (Spectrum Technologies, Aurora, IL, USA). We selected this depth because the substrate becomes very rocky at deeper depths. Upon inserting the probe vertically into the substrate, we allowed the instrument to equilibrate for a few seconds prior to measurement. We used the same calibration for all measurements. Measurements were taken at the edge of 138 individuals of 11 species: *A. lettermanii* (n=8), *A. glauca* (*n*=9), *E. congesta* (*n*=6), *Chaenactis douglasii* (Asteraceae, *n*=7), *E. lanceolatus* (*n*=17), *E. umbellatum* (*n*=16), *H. villosa* (*n*=14), *I. gordonii* (*n*=19), *L. argenteus* (*n*=21), *P. hastata* (*n*=9), and *S. crassulus* (*n*=8). For comparison, a paired measurement was also taken in a non-vegetated area at least 10 cm from each focal individual, and as far as 50 cm to avoid other plants. Soil moisture was measured after rains on 6 August 2016. This timing allowed soil conditions to equilibrate across the site and for microsite variation in soil moisture to develop over least two full days of zero precipitation.

#### Vital rates

To address Q2, we mapped plant locations and determined survival, growth, and fecundity using the following methods for each individual and census year. **Survival:** we considered plants to be alive if they produced aboveground biomass. Because vegetative dormancy is common in this community, especially in response to stress, we did not consider plants to be dead unless they failed to produce any aboveground biomass for two consecutive years. **Growth:** we calculated growth as the change in the maximum length of the individual (cm) from year *t* to year *t*+1. **Fecundity:** we counted the number of inflorescences per individual during the peak growing season.

#### Macroclimate

To capture interannual differences in macroclimate (*i*.*e*. regional climate) as a predictor in the vital rate models (Q2), we calculated the total precipitation during the growing season of each year (defined as the period between the first snow-free day in early summer and the last day before the first severe freeze in fall (below 25°F/-3.89°C), following Inouye *et al*. (2000). We included growing season precipitation in the vital rate models to capture interannual macroclimatic variation because we observed strong variation in precipitation throughout the study period including decreases in the mean and standard deviation between 2016-2019 (Figure S1). We also anticipated that growing season precipitation would be an important driver of plant population dynamics because the substrate at our site is very shallow (∼5-10 cm) and porous. All climate data were downloaded from the ‘Schofield Pass’ SNOTEL (737) station (USDA-NRCS, downloaded Oct 24, 2019), ∼4.3 km from and ∼300 m lower than the field site.

#### Belowground spatial extent

In 2015 and 2016, we measured belowground maximum rooting diameters in an area adjacent to our field site by excavating 3-5 individuals for each of 16 species and measuring maximum horizontal rooting extent (Blonder *et al*. 2018). Of the 18 species in our census plots, we were unable to measure belowground maximum diameters for two species: *Poa stenantha* and one species recorded as a single unknown seedling in 2014. To estimate the belowground spatial extent for all individuals, we calculated a taxon-specific allometric scaling factor between aboveground maximum diameter and belowground maximum diameter. For each taxon, we fitted linear regression models between aboveground and belowground maximum extent (Figure S2). For all regressions, the y-intercept was set to 0 to match the expectation of zero belowground root extent for individuals with aboveground diameters of 0 cm, recognizing that this may vary for individuals experiencing aboveground dieback. We used the product of the above-to-belowground scaling factor (*i*.*e*. the corresponding taxon-specific regression slope) and the maximum aboveground length to estimate the belowground size of all individuals. In subsequent analyses, the above-to-belowground scaling factor of *P. stenantha* was assumed to be equivalent to *E. lanceolatus* as they are both in the same subfamily (Pooideae). The calculated R^2^ values ranged from 56% (*L. argenteus*) to 99.6% (*Penstemon whippleanus*) with a mean of 88% (*sd* = 13%, *n* = 16).

#### Intraspecific and interspecific percent overlap

We measured how vegetative overlap affects vital rates using interspecific and intraspecific percent overlap as proxies for microenvironment modification. This proxy was chosen because results from the above measurements demonstrate that overlap influences microenvironment modification. However, we used overlap instead of the direct microenvironment modification values measured above, as modification may not scale additively in multi-individual and multi-species-level contexts. We defined percent overlap as the percent cumulative overlap, summed for each instance of overlap (Figures S3 & S4). To determine which individuals have overlapping distributions and the extent of overlap, we mapped each individual at our site using x-y coordinates measured in the field. We then assumed that each plant was a circular polygon with a diameter equal to its maximum length as measured in the field. Although *E. congesta, E. umbellatum*, and *H. villosa* have a greater tendency to deviate from a circular growth form than other species, especially due to dieback, this growth pattern is uncommon, and a circular polygon is appropriate in the vast majority of cases. From the mapped polygons, we determined the area of overlap for each occurrence and what percentage of each plant was overlapped by inter- and intraspecific individuals. Because plants may extend outside of the censused plot area where we do not have data on plant distributions, only the area of each plant within the plot was used in these calculations. To compare how microenvironment modification estimates vary above- and belowground, inter- and intraspecific percent overlap was similarly determined using polygons calculated based on observed aboveground extent and estimated belowground extent.

## Statistical Analyses

### Microenvironment modification by taxon (Q1)

#### Surface temperature

Because surface temperature measurements in non-vegetated areas in 2018 were paired with measurements made within individual plants, we first tested for species-level differences in minimum and maximum temperatures in 2018 using ANOVA (α=0.05) to determine whether they could be treated as comparable to the 2016 non-vegetated plot corner measurements. Minimum and maximum temperatures were the lowest and highest values recorded while deployed for each temperature data logger, respectively. We used Tukey’s honest significance test for post hoc analysis following ANOVA (α=0.05). Temperatures measured in non-vegetated areas in 2016 (at plot corners) and 2018 (10 cm from focal plants) were used as references to assess temperature modification extent and were treated similarly in all subsequent analyses.

For each year (2016 & 2018), we separately tested whether minimum and maximum surface temperatures differed between vegetated and non-vegetated areas using generalized linear models (GLM) with Gaussian families and identity link functions. As the 2016 surface temperature data was unpaired to a specific individual or taxon, we used taxon as our predictor variable and treated non-vegetated areas as an additional ‘species’ in our model. For each species, we used Tukey contrasts, α=0.05, to assess differences from non-vegetated areas. We also compared site-wide variation in minimum and maximum temperatures between vegetated and non-vegetated microenvironments for 2016 and 2018 data. For this, we used Mann-Whitney *U* tests for non-normally distributed data and Welch’s *t* tests for normally distributed data with unequal variance, as appropriate.

#### Soil moisture

We compared soil volumetric water content (%) at the edge of plants (distance = 0 cm) and paired non-vegetated areas across 11 species using a generalized linear mixed model (GLMM) with taxon, distance, and their interaction as fixed effects (family=Gaussian, link=identity). Plot was a random effect to address across-site variation. To compare soil moisture in vegetated and non-vegetated areas within taxa we used least-squares means (α=0.05). We assessed site-level variation in soil moisture between plants and paired non-vegetated areas using a Welch’s two-sample *t* test. Soil moisture data were log-transformed before analysis to achieve normality. For the site-level analysis, we also included data from 10 individuals from 5 taxa that had insufficient replication for inclusion in the taxonomic analysis: *Arnica latifolia* (*n*=4), *C. siccata* (*n*=1), *P. stenantha* (*n*=2), *Pseudocymopterus montanus* (*n*=2), and *Viola praemorsa* (*n*=1).

#### Habitat association

As larger plants are expected to have a greater capacity to modify microenvironments due to greater total biomass, the occurrence of larger plants in sites with more moderate microclimates (*e*.*g*. sites with higher soil moisture and buffered temperatures) would support the occurrence of habitat association rather than microclimate modification. In contrast, a weak relationship between plant size and environmental conditions would indicate that large individuals do not differ from small individuals in habitat, and would support that microclimate differences are due to modification by plants. To address whether microclimate differences between vegetated and non-vegetated microsites captured microclimate modification rather than fine-scale habitat association, we used estimated environmental data at 10 cm resolution to construct six GLMs (family=Gaussian). In these models, environmental variables and plant length were continuous numeric variables, and taxon and plot were factors: The structure of the GLMs was as follows:

#### Environmental variable∼Plant length*Taxon + Plot

In these models, we used six environmental variables (soil organic matter (%), soil pH, soil penetration energy (a metric of soil hardness, MJ m^-3^), median surface temperature (°C), soil moisture at 10 cm depth (g g-1), and soil moisture at 4 cm depth (g g-1)), which we extracted from a kriged raster of environmental data for the site (Dryad: https://doi.org/10.5061/dryad.33410). These microclimate data were collected in 2015-2016 at 2 m resolution and interpolated to 10 cm resolution (for full details on these data, see Blonder *et al*. (2018)). The selected environmental variables were chosen among the 27 available in this data set due to the similarity of three of the variables to the microclimate variables in this study (median surface temperature, soil moisture at 10 cm depth, and soil moisture at 4 cm depth), and because each variable captured a condition of the microhabitat that could be modified by vegetation. For these analyses, we used plant size and location data from 2015 for eleven taxa as total plant abundance was highest this year and because 2015 was when many of these environmental variables were collected. In 2015, the selected taxa (*A. glauca, C. siccata, C. douglasii, E. lanceolatus, E. congesta, E. umbellatum, H. villosa, I. gordonii, L. argenteus, P. hastata, S. crassulus*) had a maximum abundance of 624 (*L. argenteus*) and a minimum abundance of 26 (*C. douglasii*) (mean=153.7, sd=175.1). Then, for each GLM, we plotted two partial dependence plots for the environmental variables—one with plant length as the predictor variable and all other model predictors static, and the other with both plant length and taxon as model predictors and all other predictors static.

### 2) Vital rates response (Q2)

For each vital rate, we fitted an aboveground model (AG) and a belowground model (BG), using above-and belowground spatial overlap estimates. Correlation among fixed effects was low in both aboveground and belowground models (Figure S8) We used GLMMs to model vital rate responses to microenvironment modification via our vegetative overlap proxies. For each vital rate (survival, growth, fecundity) we constructed a GLMM with the following structure (in *lme4* syntax).

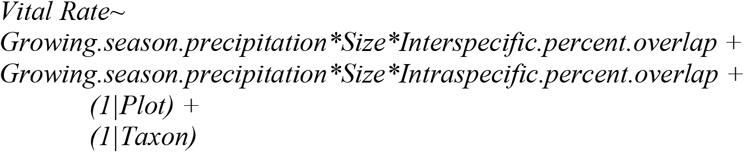

In addition to growing season precipitation and inter- and intraspecific percent overlap, plant size was included as a fixed effect because it is a strong driver of variation in vital rates, including alpine systems (Kirkpatrick 1984; Oldfather & Ackerly 2019). In all models, we used taxon-scaled size values (divided by the 90^th^ percentile across all years) to reduce variation in size across species. All fixed effects were z-transformed (zero mean, unit variance) before model fitting to allow standardized comparison and interpretation of the estimated effects. For those vital rates quantified as probabilities (survival and flowering probability), we used odds ratios to determine significance. Predictors with odds ratios <1 have a negative association with the vital rate, predictors with odds ratios >1 have a positive association, and predictors with an odds ratio of 1 have no association. We included taxon as a random effect to account for nesting of individuals within species and to estimate among-species variance in mean vital rates. Plot was included as a random effect to partially account for spatial autocorrelation. We conducted vital rate analyses for the six most abundant species (*C. siccata, E. lanceolatus, E. umbellatum, H. villosa, I. gordonii, L. argenteus*). In the growth models, the number of individuals per species ranged from 27 (*H. villosa* in 2018) to 372 (*L. argenteus* in 2016). The average number of individuals per taxon and year in the growth models was 124.6 (sd=82.7).

As required by the model structure and to address our questions, we formatted the vital rate data and constructed the GLMMs as follows.

#### Survival

we used macroclimate (growing season precipitation), size, and neighborhood data in year *t* to predict survival in year *t*+1. This model structure allowed us to include size as a predictor because by definition all plants with size=0 cm for two consecutive years are dead. We fitted a logistic regression model (binomial error distribution, logit link function). Data from the 2020 census were used to confirm the mortality of plants with size=0 cm in 2019.

#### Growth

We analyzed growth using a Gaussian model (identity link function). We used taxon-scaled growth values (divided by the 90^th^ percentile across 2015-2019) and excluded dead plants from the model because any change in their size would reflect mortality. We also excluded first-year plants and plants from the initial census year (2014) as they did not have growth data.

#### Fecundity

we analyzed fecundity using two models. For the first model, we fitted a logistic regression model (binomial error distribution, logit link function) to model flowering probability. For the second model, we analyzed the number of inflorescences of flowering individuals using a Gaussian model (identity link function). Dead individuals and seedlings were excluded from both fecundity models, while non-flowering individuals were excluded from the model of the number of inflorescences. To account for taxonomic differences in inflorescence number, the number of inflorescences was scaled by dividing by the 90^th^ fecundity percentile across all years for each taxon because the number of inflorescences is comparable within, but not between species. The scaled fecundity values were also log-transformed to limit overdispersion. Model residuals met assumptions in all models.

We conducted all analyses in R, version 4.1.2 (R Core Team 2021). We used *lme4* (Bates *et al*. 2021) to calculate the GLMMs. Post-hoc tests for the microenvironment analyses were done using *multcomp* (Hothorn *et al*. 2022) and *emmeans* (Lenth *et al*. 2022). We used the packages *sf* (Pebesma *et al*. 2021a), *sp* (Pebesma *et al*. 2021b), and *rgeos* (Bivand *et al*. 2021) to estimate spatial overlap. We used *DHARMa* (Hartig & Lohse 2021) to check for over- and underdispersion of the residuals and *PerformanceAnalytics* (Peterson *et al*. 2020) to assess correlations among model terms. *GGally* (Schloerke *et al*. 2021) was used to generate correlation plots and we used *pdp* (Greenwell 2017) to generate partial dependence plots.

## Results

### Microenvironment modification by taxon

#### Surface temperature

In non-vegetated areas in 2018, maximum temperatures ranged from 51.5-64.0 °C across species (Figure S5a). We detected no significant differences in maximum temperatures among species (ANOVA: F_6, 73_ = 1.75, P =0.12). In contrast, the range of minimum temperatures in non-vegetated areas in 2018 was much narrower (1.5-4.5 °C) and differed depending on the focal plant species (ANOVA: F_6, 73_ = 2.23, P =0.05), likely reflecting species-specific variation in microenvironment preference (Figure S5b). However, we did not detect differences in minimum temperatures in post hoc pairwise comparisons of species, though differences between *L. argenteus* and *A. congesta* were close to the α=0.05 threshold (Tukey contrast: P=0.054).

Maximum temperatures were lower within versus outside vegetation in 2016 (Mann-Whitney: U = 1532, n_1_ = 91, n_2_ = 83, P < 0.001) and 2018 (Welch’s two-sample t-test: t=-5.1572, *v* =122.54, P < 0.001) (Figure S6). Maximum surface temperatures were lower within the canopies of 7 out of 9 species in 2016 and 4 out of 7 species in 2018 compared to non-vegetated areas (Figure 2a). We observed cooling effects for all focal erect monocotyledons: *A. lettermanii* (Mean cooling, °C: 2016=-6.80); *E. lanceolatus* (2016=-5.19; 2018=-4.40). Among erect dicotyledons (*A. glauca, L. argenteus, S crassulus*), we only observed cooling effects for *L. argenteus* (2016=-5.58, 2018=-8.02). For rosette dicotyledons, maximum temperatures were lower in *I. gordonii* (2016=-6.30, 2018=-4.15) and *P. hastata* (2016=-8.73), but not *E. congesta*. Finally, for deciduous dicotyledonous shrubs, maximum temperatures were cooler for *H. villosa* in both years (2016=-4.97, 2018=-6.23), but were only cooler for *E. umbellatum* in 2016 (−6.17).

**Figure 2:**
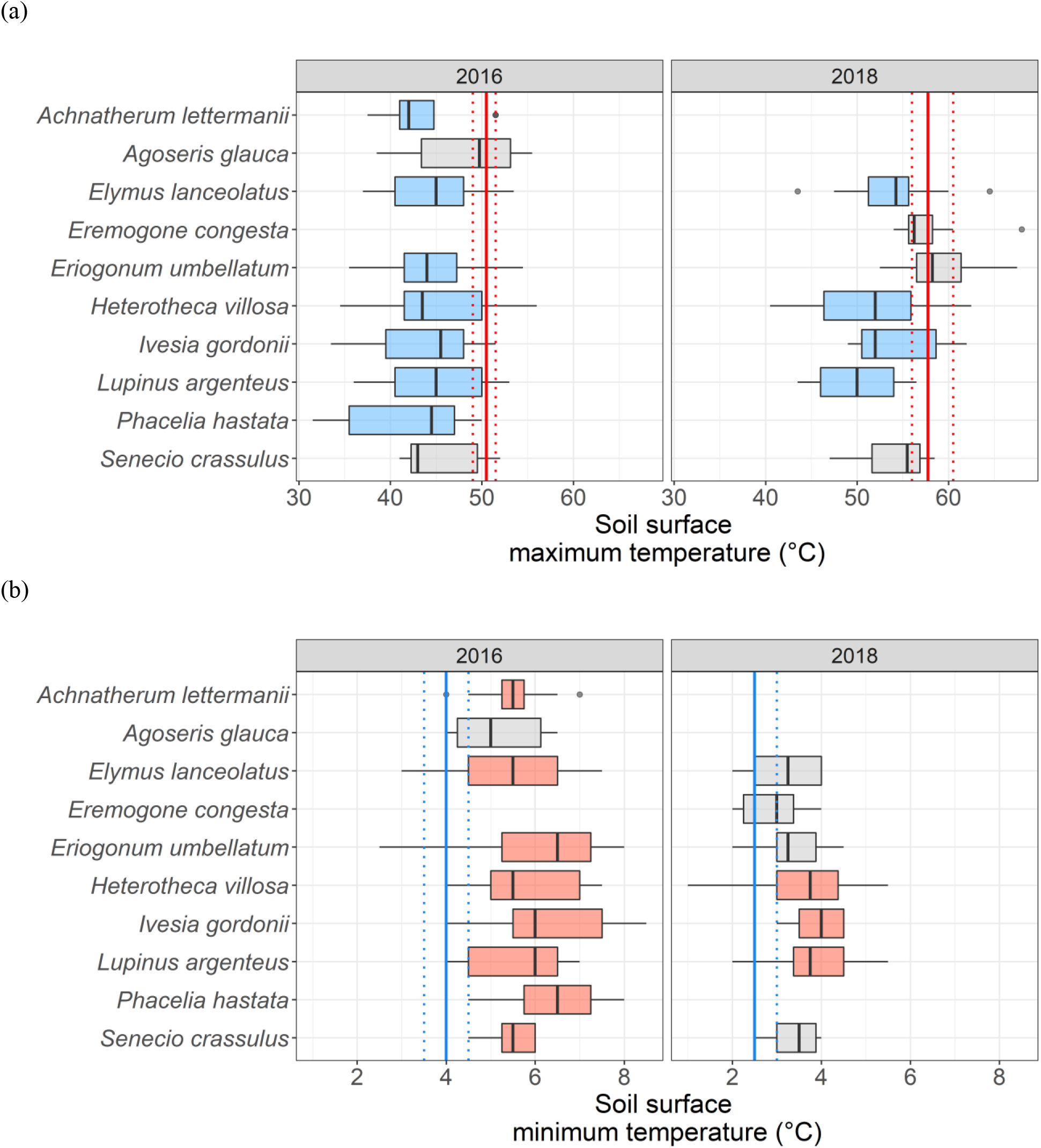
(a) Maximum and (b) minimum temperatures (°C) of data loggers within the canopies of selected alpine plant species (*n* = 9 in 2016 and 7 in 2018) and non-vegetated areas. The selected taxa were among the most abundant in the community and represented a range of growth forms. All focal plants had a vegetative diameter > 1 cm. Vertical bands indicate measured values in non-vegetated areas: leftmost line=1^st^ quartile, middle line=median, rightmost line=3^rd^ quartile, with red bands for maximum temperatures and blue bands for minimum temperatures. For minimum temperatures in 2018, the first quartile is equal to the median. Blue and red boxplots indicate statistically significant differences from non-vegetated areas.

Minimum temperatures were higher within versus outside vegetation in 2016 (Mann-Whitney: U = 6678, n_1_ = 91, n_2_ = 83, P < 0.001) and in 2018 (Mann-Whitney: U = 4940.5, n_1_ = 80, n_2_ = 80, P < 0.001) (Figure S6). Minimum temperatures were higher within the canopies of 8 out of 9 species than in non-vegetated areas in 2016 and 3 out of 7 species in 2018 (Figure 2b). Minimum temperatures were warmer for erect monocotyledons in 2016 (mean warming, °C: *A. lettermanii* (2016=+1.51); *E. lanceolatus* (2016=+1.43), but not for *E. lanceolatus* in 2018. Warmer minimum temperatures were also observed for some, but not all, rosette dicotyledons and years: *L. argenteus* (2016=+1.73, 2018=+1.13); *S. crassulus* (2016=+1.51). Modification trends for warming for rosette dicotyledons and deciduous dicotyledonous shrubs were similar to those found for cooling. Warming was observed in 2016 and 2018 in *I. gordonii* (2016=+2.31, 2018=+1.26), *P. hastata* (2016=+2.43), and *H. villosa* (2016=+1.84, 2018=+0.90), but was only observed in 2016 for *E. umbellatum* (+2.23).

#### Soil moisture

At the site level, soils were wetter within plants compared to non-vegetated areas (Welch’s two-sample t-test: t= 6.2668, *v* =271.97, P < 0.001) (Figure S7). The substrate below 5 out of 11 plant species had higher volumetric water content (%) than in adjacent non-vegetated areas (mean difference below plants, %: *E. umbellatum* (+1.87), *I. gordonii* (+3.33), *L. argenteus* (+1.99), *P. hastata* (+2.48), and *S. crassulus* (+5.34)) (Figure 3). Two additional species (*A. lettermanii and H. villosa*) trended towards wetter soil at the plant level (median in plant values > 3^rd^ quartile non-vegetated values), but with weak statistical support.

**Figure 3:**
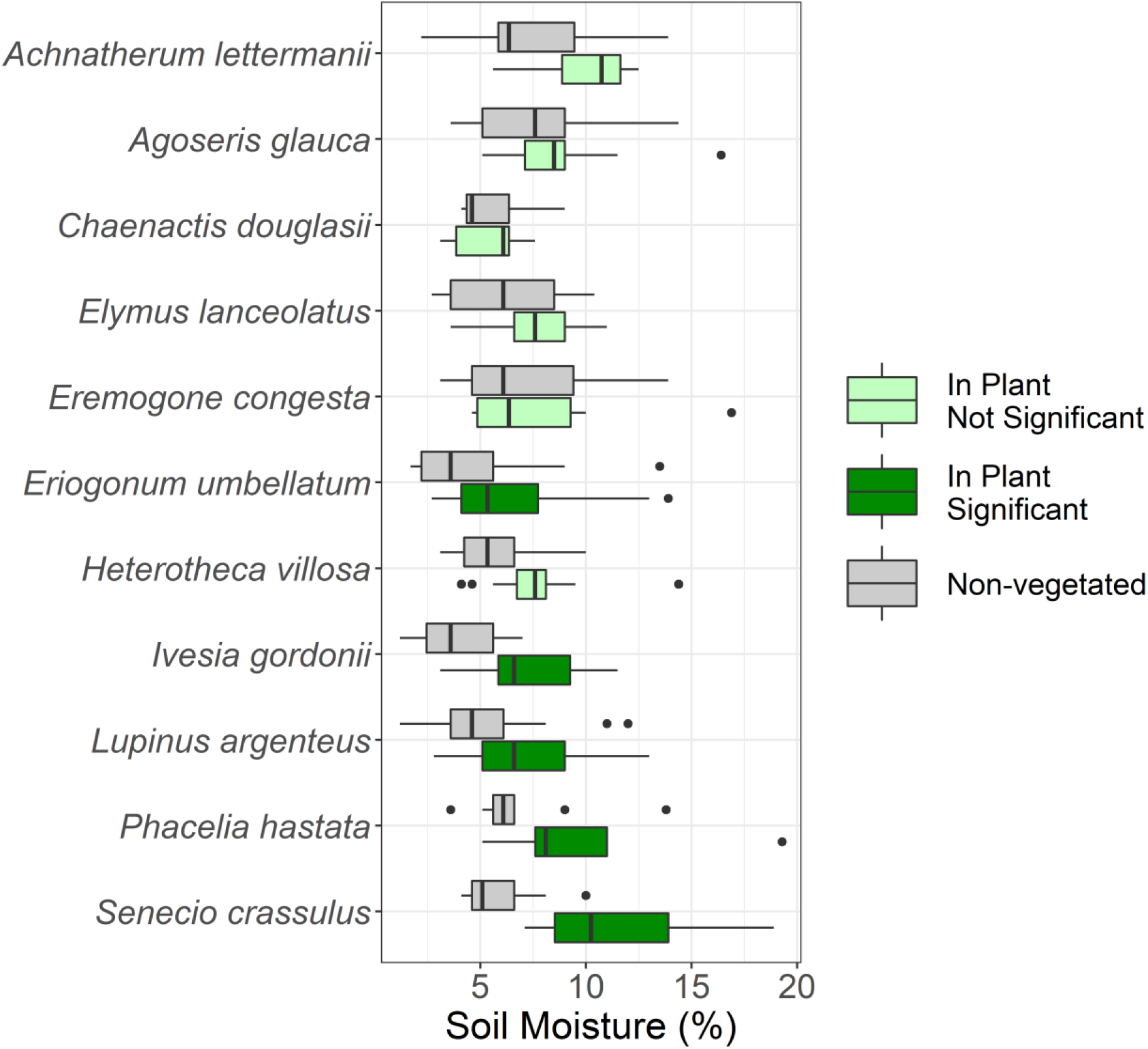
Soil moisture inside and outside of plants for 11 species in 2016. The selected taxa were among the most abundant in the community and represented a range of growth forms. All focal plants had a vegetative diameter > 1 cm. Dark green boxes indicate statistically significant differences compared to the non-vegetated comparison.

#### Habitat association

We observed positive dependencies of soil organic matter, soil moisture at 10 cm depth, and soil moisture at 4 cm depth on plant length and negative dependencies of soil pH, soil penetration energy, and median surface temperature on plant length (Figure S9), with high variation among taxa (Figure S10). However, the magnitude of the response scale was small for all environmental variables.

### 2) Vital rate responses

For our microclimate modification proxies, interspecific and intraspecific overlap, effects on vital rates varied. Interspecific percent overlap had negative effects on survival probability (AG: 0.83–1.00; BG: 0.65–0.81), flowering probability (AG: 0.75–0.94; BG: 0.68–0.89), and number of inflorescences (AG: -0.13–-0.02), but a positive effect on growth (BG: 0.05–0.13). Intraspecific percent overlap, on the other hand, had a positive effect on flowering probability (BG: 1.03–1.35) and a negative effect on inflorescence production (BG: -0.17–-0.04).

The effects of species interactions on vital rates were also context dependent. For larger plants, interspecific percent overlap had a weaker effect on growth (AG: -0.08–-0.01), flowering probability (BG: 0.72–0.88), and inflorescence production (AG: -0.10–0.00; BG: -0.13–-0.04). In contrast, for larger plants, there was an increased positive effect of intraspecific overlap on survival (AG: 1.15–1.39; BG: 1.47–1.82), growth (AG: 0.01–0.08), flowering probability (BG: 1.34–1.78), and inflorescence production (BG: 0.02–0.12).

The effects of species interactions on vital rates also depended on interactions with macroclimate. In wetter years, there was a positive effect of interspecific overlap on survival (BG: 1.02–1.18) and growth (AG: 0.04–0.10; BG: 0.01–0.07). Flower production was lower for plants with greater interspecific overlap in wetter years (AG: -0.11–-0.01; BG: -0.11–0.00). However, intraspecific percent overlap, had a positive effect on inflorescence production in wetter years (BG: 0.00–0.11). Above- and belowground model results for survival, growth, flowering probability, and number of inflorescences are in Figures 4-7, with numeric model results in Tables S1*-*S8. Parameter estimates for the random effects for all models are in Figures S11-S18.

**Figure 4:**
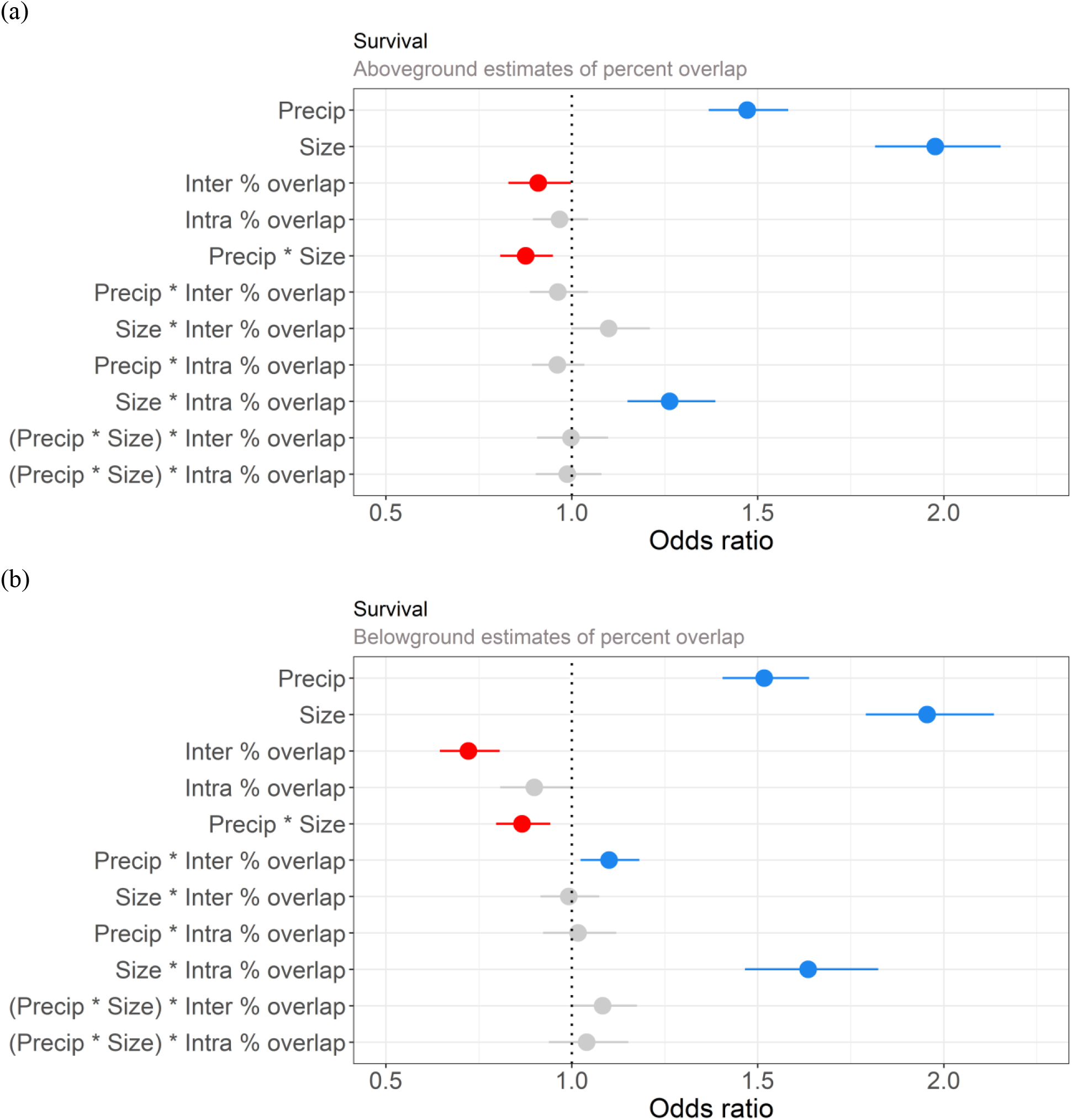
Odds ratios with 95% confidence intervals for standardized fixed effects in the models describing survival probability. Estimates of vegetative percent overlap were based on (a) aboveground and (b) belowground biomass.

**Figure 5:**
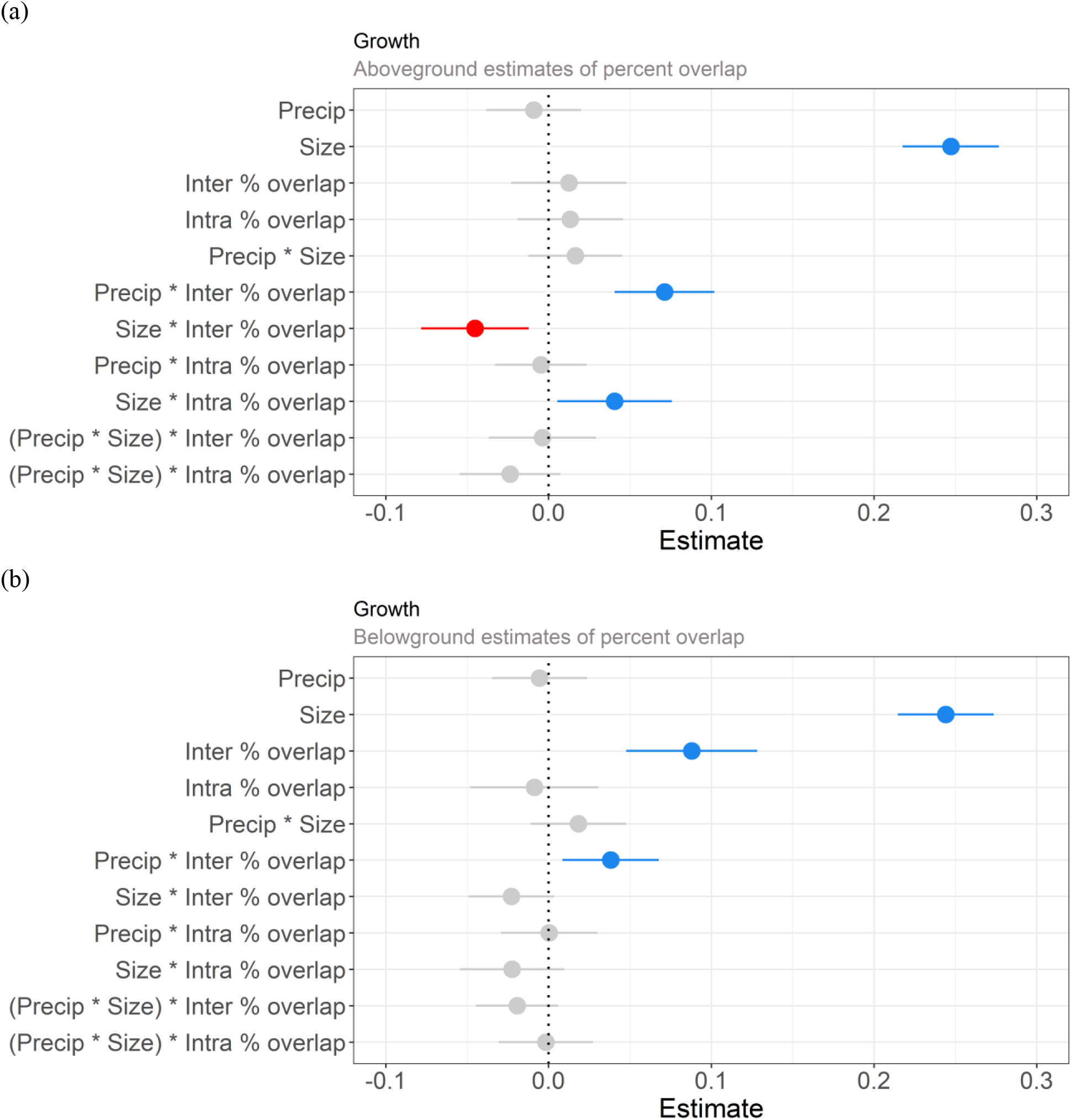
Parameter estimates with 95% confidence intervals for standardized fixed effects in the models describing growth. Estimates of vegetative percent overlap were based on (a) aboveground and (b) belowground biomass.

**Figure 6:**
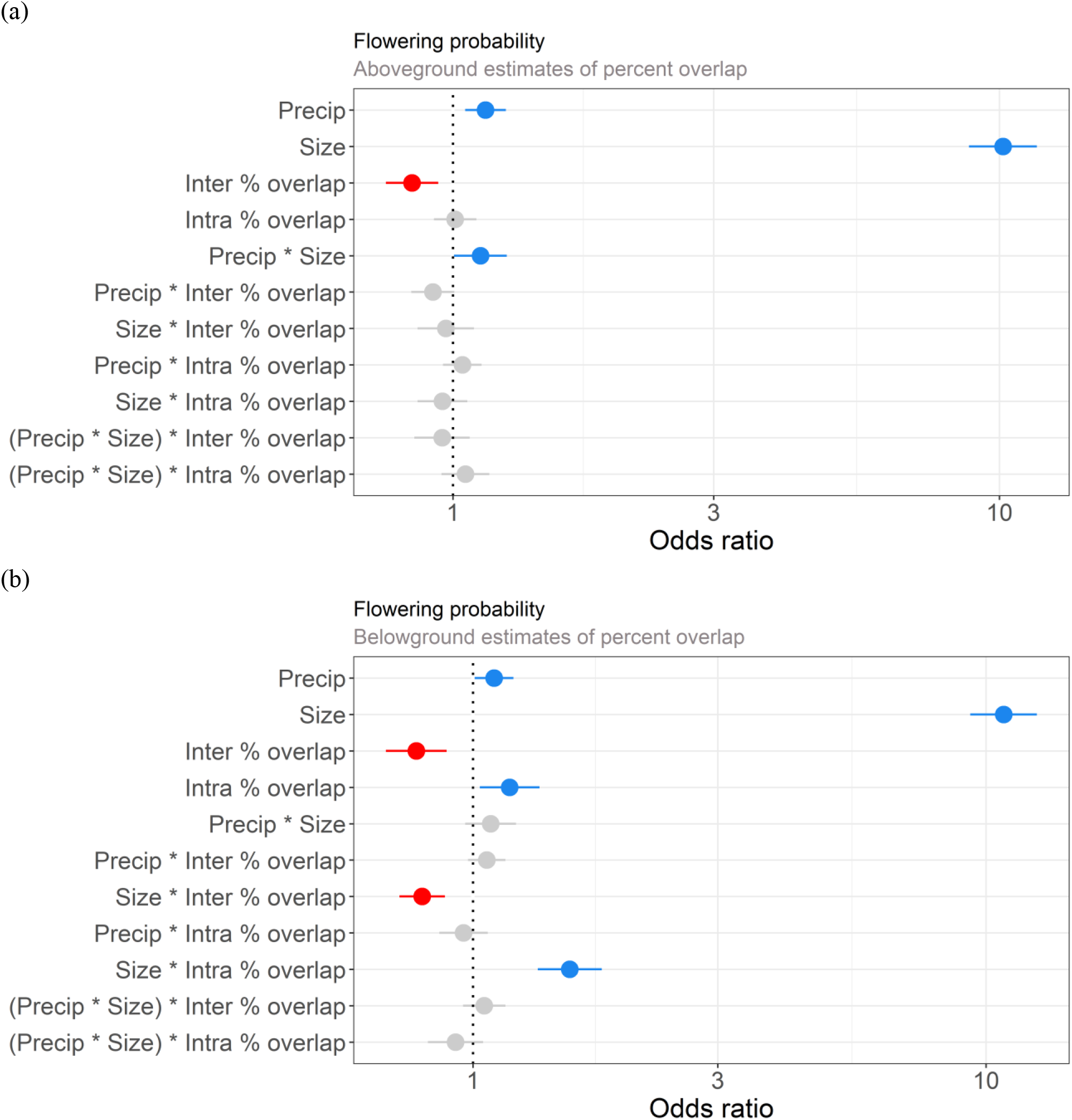
Odds ratios with 95% confidence intervals for standardized fixed effects in the models describing flowering probability. Estimates of vegetative percent overlap were based on (a) aboveground and (b) belowground biomass. The x-axes were log_10_ transformed to improve data visualization.

**Figure 7:**
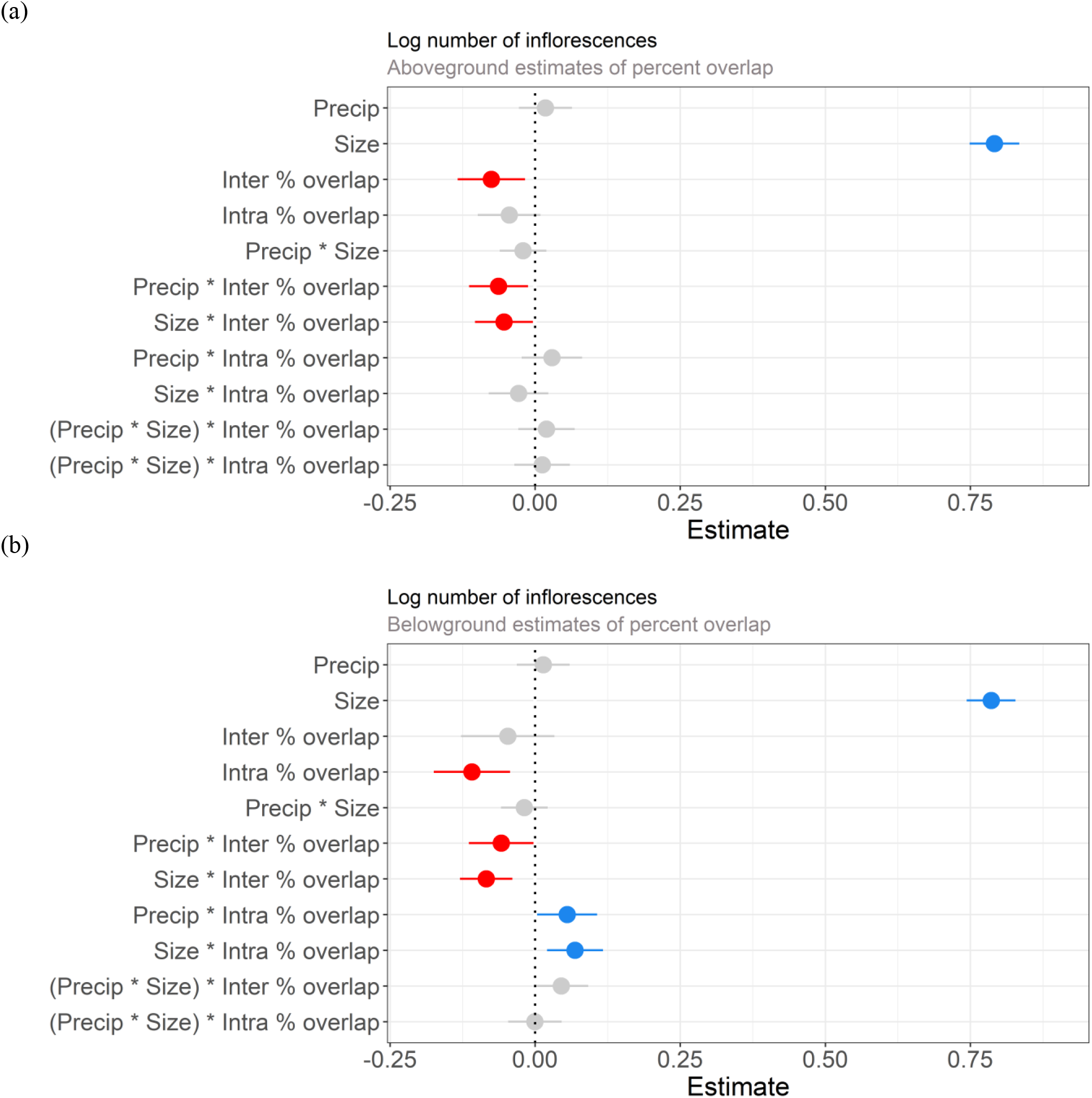
Parameter estimates with 95% confidence intervals for standardized fixed effects in the models describing the log number of inflorescences. Estimates of vegetative percent overlap were based on (a) aboveground and (b) belowground biomass.

Additionally, we observed positive effects of growing season precipitation and size on vital rates. Survival increased in years with higher precipitation (Aboveground model (AG): confidence interval=1.37–1.58; Belowground model (BG): 1.41–1.64). Flowering was also more likely in wetter years (AG: 1.05–1.25; BG: 1.01–1.20). Larger plants survived better (AG: 1.82–2.15; BG: 1.79–2.13) and grew more (AG: 0.22–0.28; BG: 0.21–0.27). Flowering was more likely for larger plants (AG: 8.79–11.71; BG: 9.32–12.57) and larger plants produced more inflorescences (AG: 0.75–0.83; BG: 0.74–0.83). Larger plants were more likely to flower in wetter years (AG: 1.01–1.26). However, the effect of size on survival was weaker in wetter years in the aboveground and belowground models (AG: 0.81–0.95; BG: 0.80–0.94).

## Discussion

In this alpine plant community, plants modified microenvironments by buffering temperature extremes (lower maximum and higher minimum surface temperatures) and by increasing soil moisture relative to open areas. Variation in multiple vital rates was attributable to variation in aboveground and belowground microenvironment modification proxies (inter- and intraspecific percent overlaps), which were frequently context dependent. These results illustrate the complex interplay between spatial overlap, microenvironment, and macroclimate on demography and community assembly.

### Microenvironment

Surface temperatures and soil moisture differed between vegetated and non-vegetated microsites. Nearly all considered taxa buffered temperatures in this community and 5 out of 11 increased soil moisture. Temperature buffering operated separately from increasing soil moisture for some taxa. For example, *A. lettermanii, E. lanceolatus*, and *H. villosa* buffered maximum and minimum temperatures in 2016 but did not increase soil moisture. On the other hand, soil moisture near *S. crassulus* was higher than non-vegetated areas in 2016, but we did not detect buffered maximum temperatures in the same year.

As alpine areas are frequently characterized by high levels of microenvironment heterogeneity, with plant distributions reflective of this variation (Rae *et al*. 2006; Opedal *et al*. 2015; Ohler *et al*. 2020), these microclimate differences may reflect plant microenvironment preference. However, we consider this to be unlikely because nearly all non-vegetated microclimate data were collected at fine-scales—10 cm from the edge of the focal plants. Also, as we observed limited dependency of six environmental variables related to microenvironment modification on plant size; thus, plants with greater capacity to modify microenvironments do not preferentially occupy more buffered microsites (Figures S9-S10). Finally, our prior work examining other abiotic gradients (Blonder *et al*. 2018) suggests it is unlikely that sub-meter scale variation or modification in soil texture or nutrient availability is occurring, though we lack data to directly assess such factors.

Our results indicate that the extent of microenvironment modification can vary across years. We observed fewer statistically detectable differences in surface temperature buffering in 2018 compared to 2016, although nearly all taxa trended towards temperature buffering for both years. Different methodology may have contributed to lower buffering values in 2018 compared to 2016 as focal plants varied between years and non-vegetated measurements were typically taken spatially closer to vegetation in 2018. Decreased plant health may have also led to weaker surface temperature buffering in 2018. Average growing season precipitation values were low in 2016-2018 (Figure S1), and after multiple drought years, plants in 2018 appeared sparser compared to previous years. Buffering temperatures may have also been more difficult in 2018 as median soil surface temperature maximums were higher in 2018 than in 2016 and median soil surface temperature minimums were lower in 2018 than in 2016.

### Context dependency of interactions

The effects of inter- and intraspecific percent overlap varied among vital rates and depended on whether above-or belowground spatial extents were considered in the models. As belowground extents were estimated from aboveground plant length, differences between model results using above-versus belowground percent cover could be greater if ‘true’ rather than estimated belowground extents could be included. However, even with estimated belowground extents, our results demonstrate the sensitivity of the vital rate analyses to the microenvironment modification proxy used and highlight the importance for future studies to consider how the spatial extents of modification may differ across ecological realms (*e*.*g*. aboveground and belowground).

Interspecific overlap had detectable effects on vital rates in six out of eight models, five of which were negative. In comparison, we found positive effects of intraspecific percent overlap on flowering probability (belowground, BG), but negative effects on the number of inflorescences produced (BG). Based on our microenvironment data and microenvironment modification results from previous studies in tundra systems (*e*.*g*. (Nystuen *et al*. 2019; Mallen-Cooper *et al*. 2021), we expect that plants with higher inter-or intraspecific percent overlap experience a more buffered microenvironment compared to plants with low to no overlap. Thus, the divergent responses in vital rates to inter- and intraspecific percent overlap might indicate greater competition among heterospecific neighbors compared to conspecific neighbors. Alternatively, this divergence may represent different demographic strategies under distinct microenvironment conditions. However, we are unable to disentangle the microenvironment effect from other possible effects of overlap, including increased structural support, and higher pollinator visitation rates. As many plants in this community grow in dense, heterogeneous clusters, individuals may simultaneously experience an array of competitive and facilitative neighbor effects that cannot be disentangled through net effect measurements like ours (Bakker *et al*. 2018).

We found that growing season precipitation was a positive driver of survival (aboveground, AG & BG) and flowering probability (AG & BG). We observed weaker effects of precipitation on survival for larger plants (AG & BG), suggesting that larger individuals were less prone to drought-induced mortality. Larger plants were also more likely to flower with increased growing season precipitation (AG), further indicating that demographic strategies may differ for large versus small individuals.

Macroclimatic conditions varied substantially during this study and species interactions often varied in response to variation in growing season precipitation. Because water limitation may have led to widespread physiological stress in this community, facilitation among species may be limited during drought years (Maestre *et al*. 2009) or masked by competitive effects. Under lower growing season precipitation, intraspecific overlap negatively affected the number of inflorescences (BG), and interspecific overlap negatively affected growth (AG & BG) and survival (BG). We also did observe some evidence for facilitation in dry years which may be due to microclimate modification—a negative interaction between precipitation and interspecific overlap on number of inflorescences (AG & BG). As the integrated effects of environmental context on vital rates determine plant performance, the contrasting effects of these predictors may represent demographic tradeoffs (Reznick 1983; Stearns 1989) with multiple potential drivers. For example, prioritizing growth and thus accessibility to resources could be advantageous when potential interspecific competitors are also benefiting from favorable environmental conditions. Additionally, there could also be facilitative effects between growing season precipitation and interspecific and intraspecific overlap if neighboring plants are better able to ameliorate the microhabitat under good conditions, such as by increasing soil water retention or shading due to higher leaf output.

By directly testing for microenvironment modification and pairing that to data on vegetative overlap, we show that modification is associated with variation in vital rates. Previously at this site, Blonder *et al*. (2018) found that spatial distributions of plants were predictable by microenvironment (including soil moisture, soil nutrients, and surface temperatures), neighborhood density, and plant functional traits. This finding was based partially on neighborhood density indices, which are less precise proxies for microenvironment modification. Additional levels of complexity influencing vital rates, but not included in this study include species-level variation in vital rate response (Jongejans & Kroon 2005) and vital rate lability (Jongejans *et al*. 2010).

### Implications for community assembly

First, our results indicate that microenvironment modification affects two community assembly processes, environmental filtering (whether a plant can survive and persist in a given habitat) and biotic interactions (with other community members) (HilleRisLambers *et al*. 2012). Second, shifts in spatial patterning due to microenvironment modification likely can result in order-dependent assembly, as seedling establishment depends on habitats created by resident species.

How microenvironments are modified by plants has important implications for how communities will respond to climate change (Anthelme *et al*. 2014). Future alpine communities may experience increased dependence on species interactions to buffer temperature extremes and decreases in water availability. However, macroclimate shifts may also lead to context-dependent effects on microclimate modification and thus on demography. For example, growing season precipitation was a positive predictor of plant survival and flowering probability in our models. In response to the predicted intensification of drought conditions in this region (Cook *et al*. 2015; Williams *et al*. 2020), this study site is likely to experience shifts in species composition and abundance. Critically, these community shifts could drive feedbacks on species interactions, including microenvironment modification, propelling further community change. Also, climate change response may vary due to the high habitat heterogeneity and decoupling from atmospheric conditions that often occur in alpine communities (Körner 2003; Scherrer & Körner 2011; Roth *et al*. 2014).

### Conclusions

In this alpine plant community, we demonstrated substantial microenvironment modification by multiple species, and that the strength of microenvironment modification can vary across years in response to variation in macroclimatic conditions. We observed positive, negative, and neutral effects of inter- and intraspecific vegetation overlap on vital rates, with frequent dependency on context (individual plant size and growing season precipitation). This study indicates that climate change may affect species’ ability to modify microenvironments, as well as the effect of microenvironment modification on vital rates.

