## Supplemental Materials for "Linking microenvironment modification to species interactions and demography in an alpine plant community"

Figure S1: Incremental precipitation (cm) from 2014-2019 for days with rain during the growing season. Growing season average per day (mm), **2014**: 2.5 ± 6.3 (sd); **2015**: 3.1 ± 5.6; **2016**: 1.3 ± 3.3; **2017**: 1.3 ± 4.2; **2018**: 1.1 ± 3.3; **2019**: 0.5 ± 1.8).

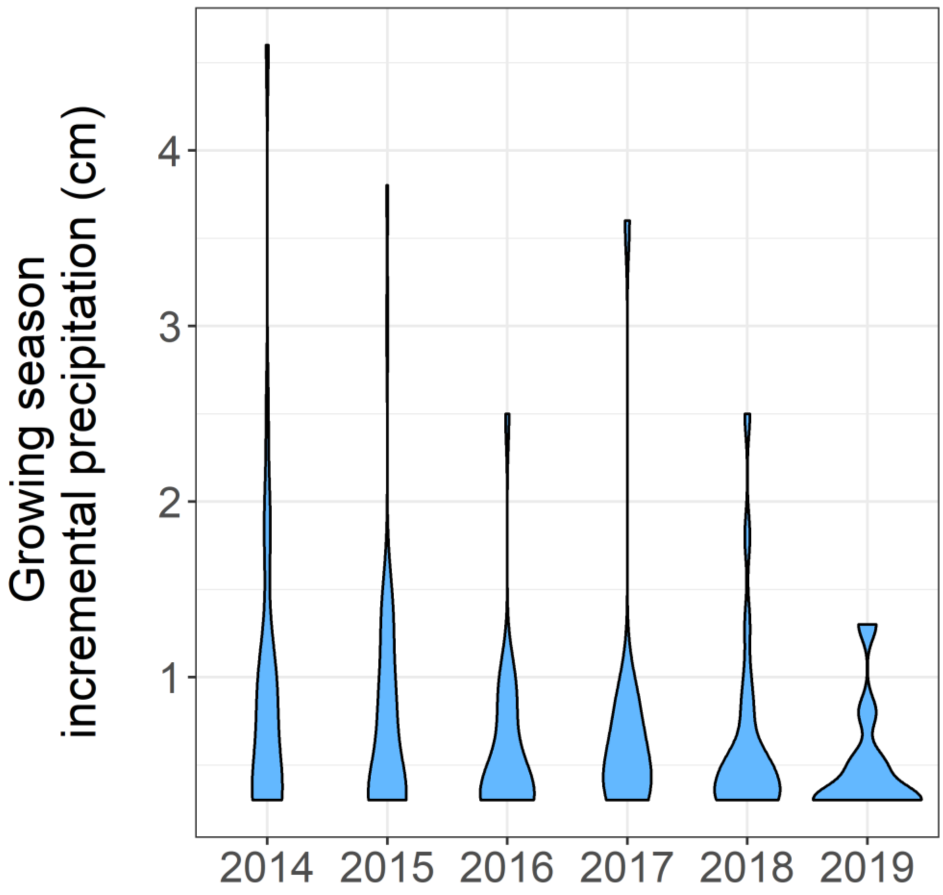

Figure S2: Linear regression between belowground and aboveground diameter for each sampled species. The regression intercept was forced through zero to account for the expectation of no belowground extent for plants with an aboveground diameter of 0.

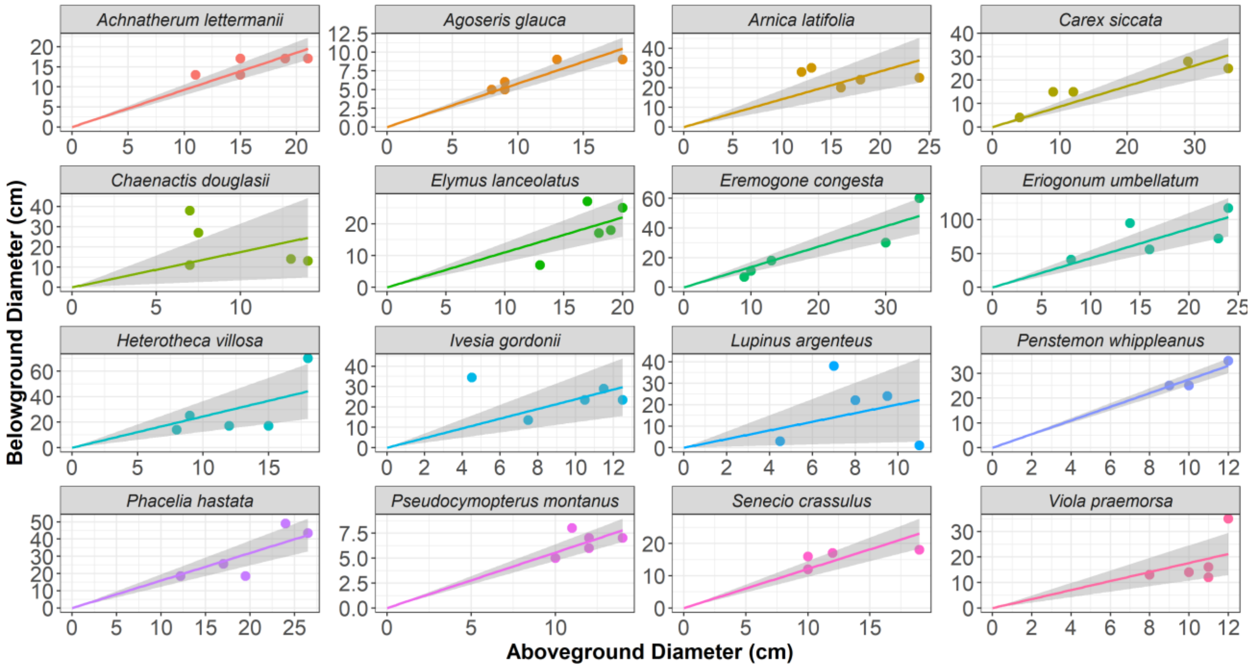

Figure S3: Schematic (a) for the determination of vegetative overlap metrics (interspecific percent overlap, intraspecific percent overlap) and their calculated values (b) for four individuals (A, B, C, D). In this example, polygons with the same color represent the same species (A & B=green, C=yellow, D=dark green).

(a)

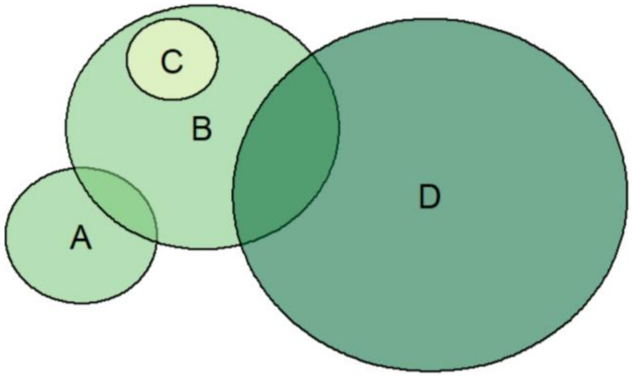


(b)

| Individual | Interspecific Percent overlap (%) | Intraspecific Percent overlap (%) |
|---|---|---|
| A | 0 | 25 |
| B | 43 | 8 |
| C | 100 | 0 |
| D | 15 | 0 |

Figure S4: Interspecific and intraspecific percent overlap counts using aboveground and belowground spatial data for all taxa in 2015. Percent overlaps range from 0% to 1200%, with each bar capturing 20% of that range.

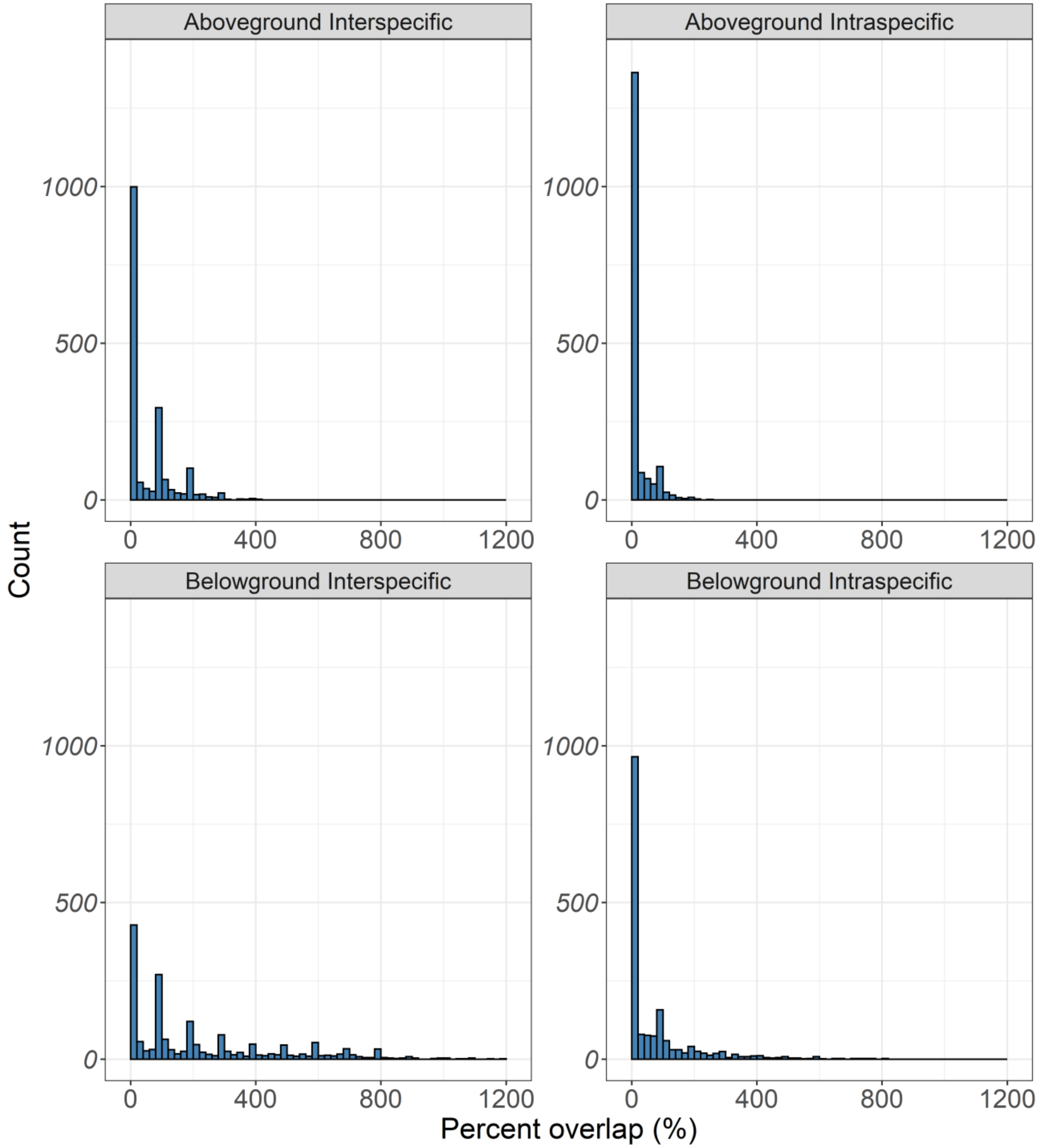

Figure S5: (a) Maximum and (b) minimum temperatures (°C) in non-vegetated areas 10 cm outside of focal taxa in 2018

(a)

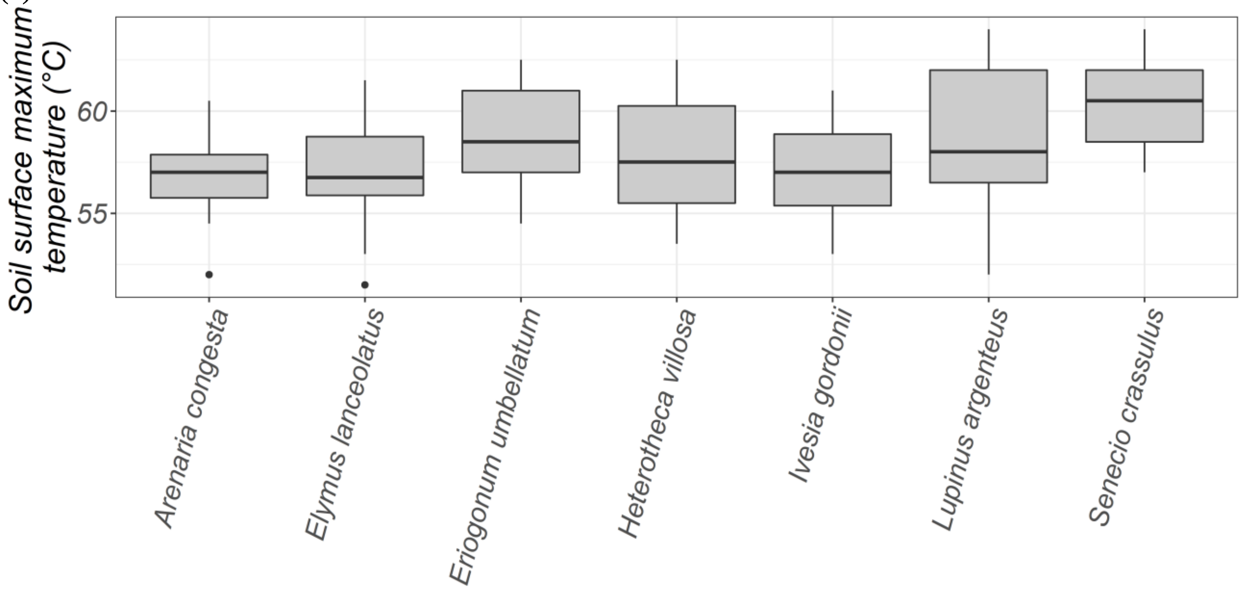


(b)

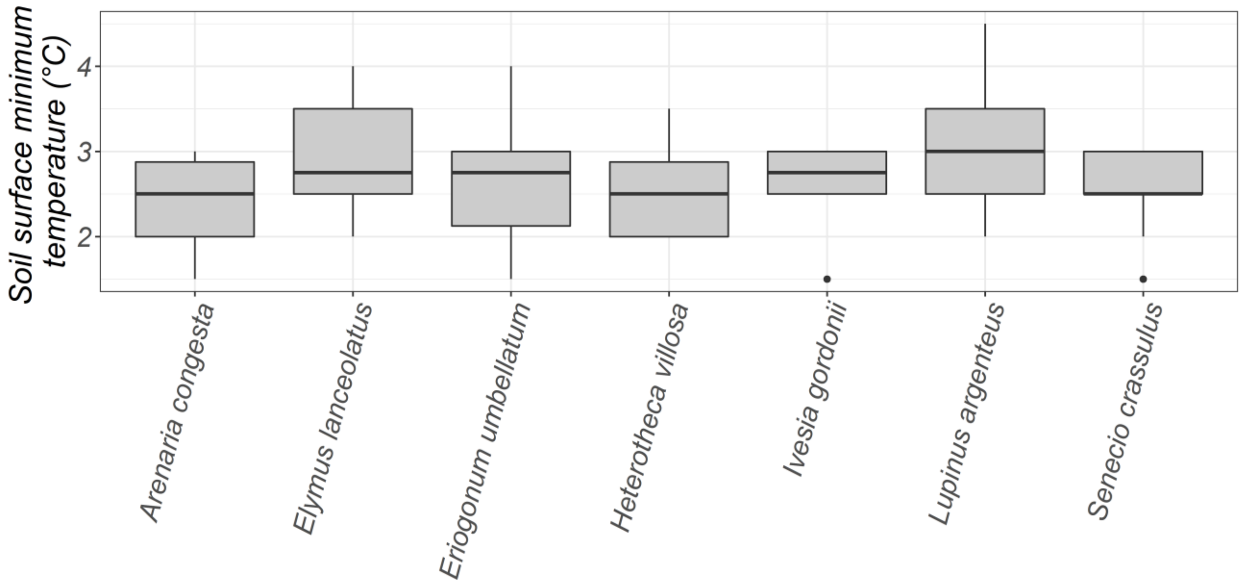

Figure S6: Temperatures (°C) in vegetated versus non-vegetated microenvironments for (a) minimum temperature 2016, (b) maximum temperature 2016, (c) minimum temperature 2018, and (d) maximum temperature 2018. For all comparisons, differences between ‘In Plant’ and ‘Non-vegetated’ temperatures were significant.

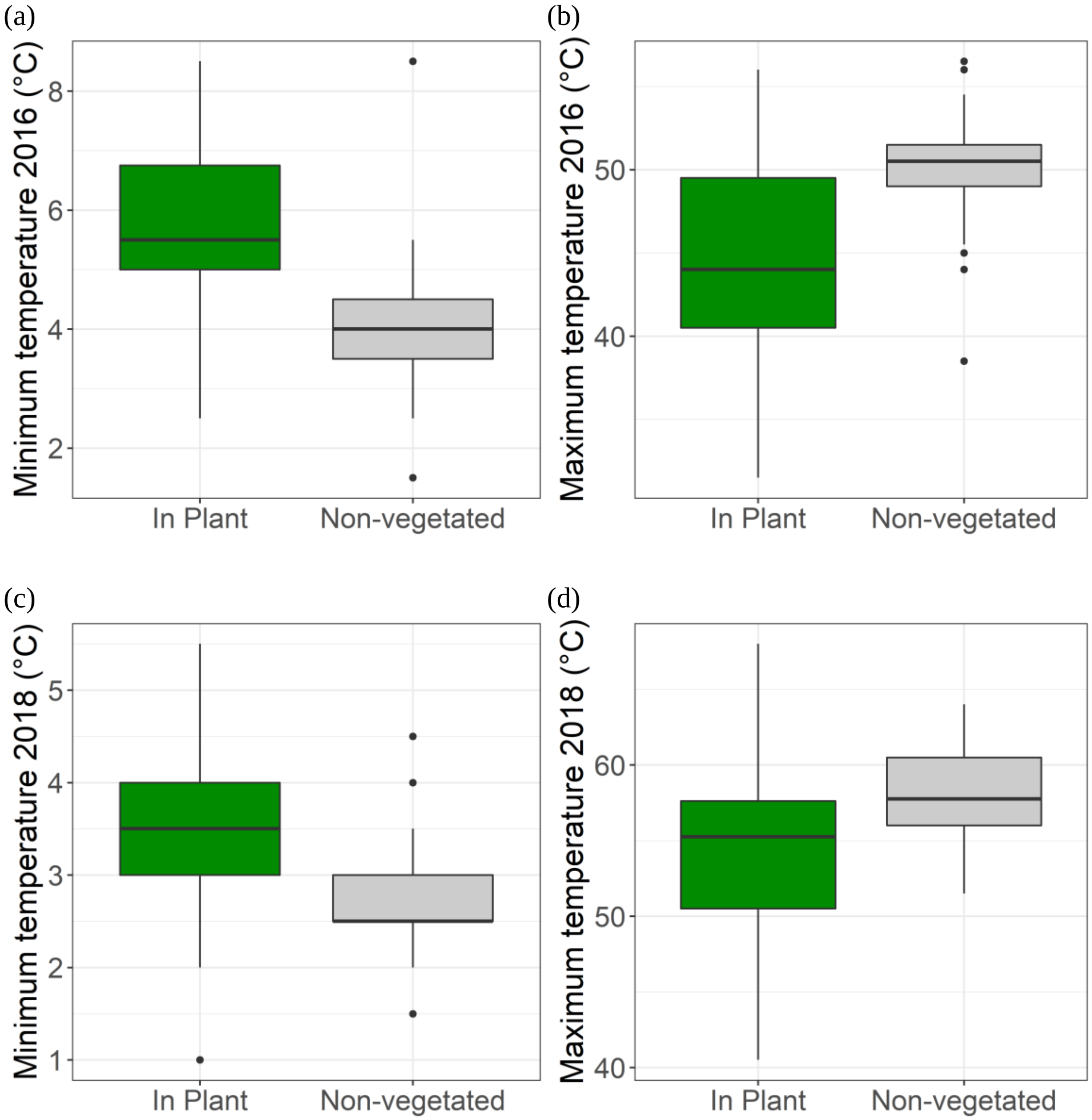

Figure S7: Log soil moisture (%) in vegetated and non-vegetated microenvironments in 2016. Differences between microenvironment types were significant at $\alpha=0.05$.

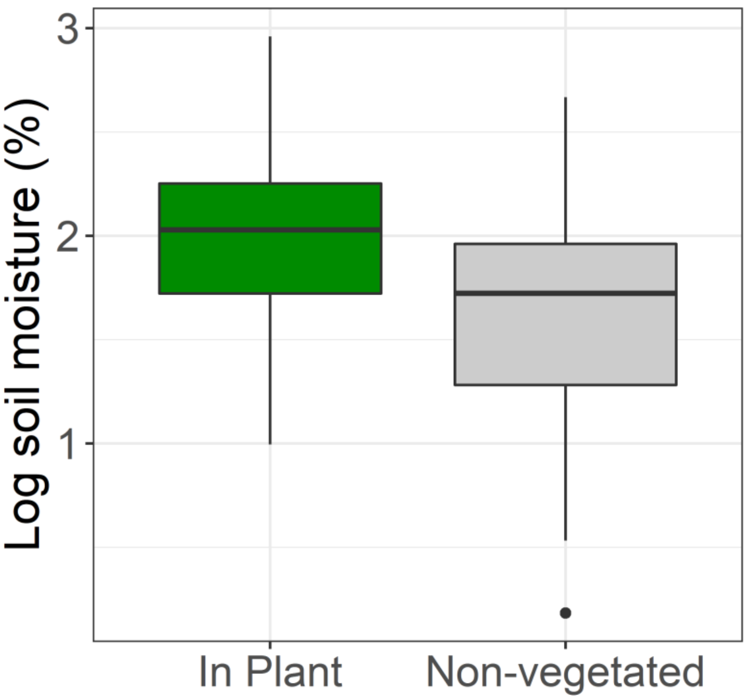

Figure S8: Correlation plots of fixed effects used in vital rate regression models of survival, growth, flowering probability, and number of inflorescences with parameters counted using (a) aboveground estimates and (b) belowground estimates of spatial overlap. Correlations were calculated as pairwise Pearson coefficients using all 6 years of demography data for *E. lanceolatus*, *E. umbellatum*, *H. villosa*, *I. gordonii*, and *L. argenteus* and did not include first-year individuals (*i.e.* seedlings and recruits).

(a) Using **aboveground** estimates of overlap

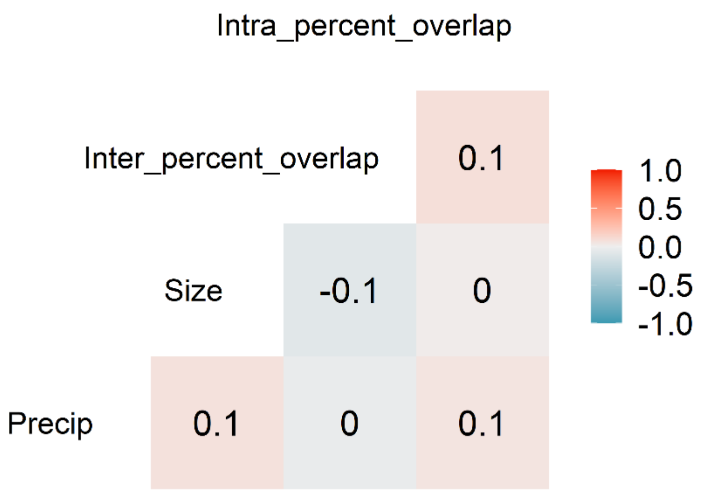


(b) Using **belowground** estimates of overlap

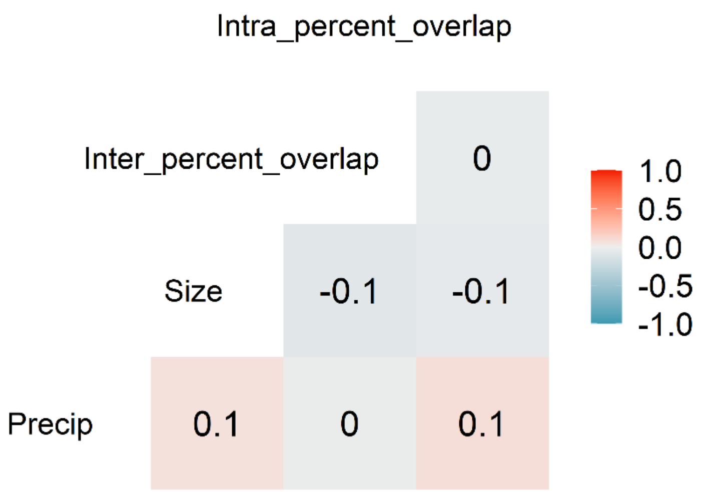

Figure S9: Partial dependence plots from six separate Generalized Linear Models (GLMs) for the response of environmental variables (median surface temperature (°C), soil pH, soil penetration energy (MJ $m^{-3}$), soil organic matter (%), soil moisture at 10 cm depth (g g-1), and soil moisture at 4 cm depth (g g-1)) to plant length (cm), taxon (n=17), plot, and the interaction between plant length and taxon. Plant length and location data were from the 2015 annual census, while environmental data for each plant location were extracted from kriged spatial data (10 cm resolution) from Blonder *et al.* (2018). Y-axis values are the average predicted values of the environmental variable based on plant length with other model predictors static.

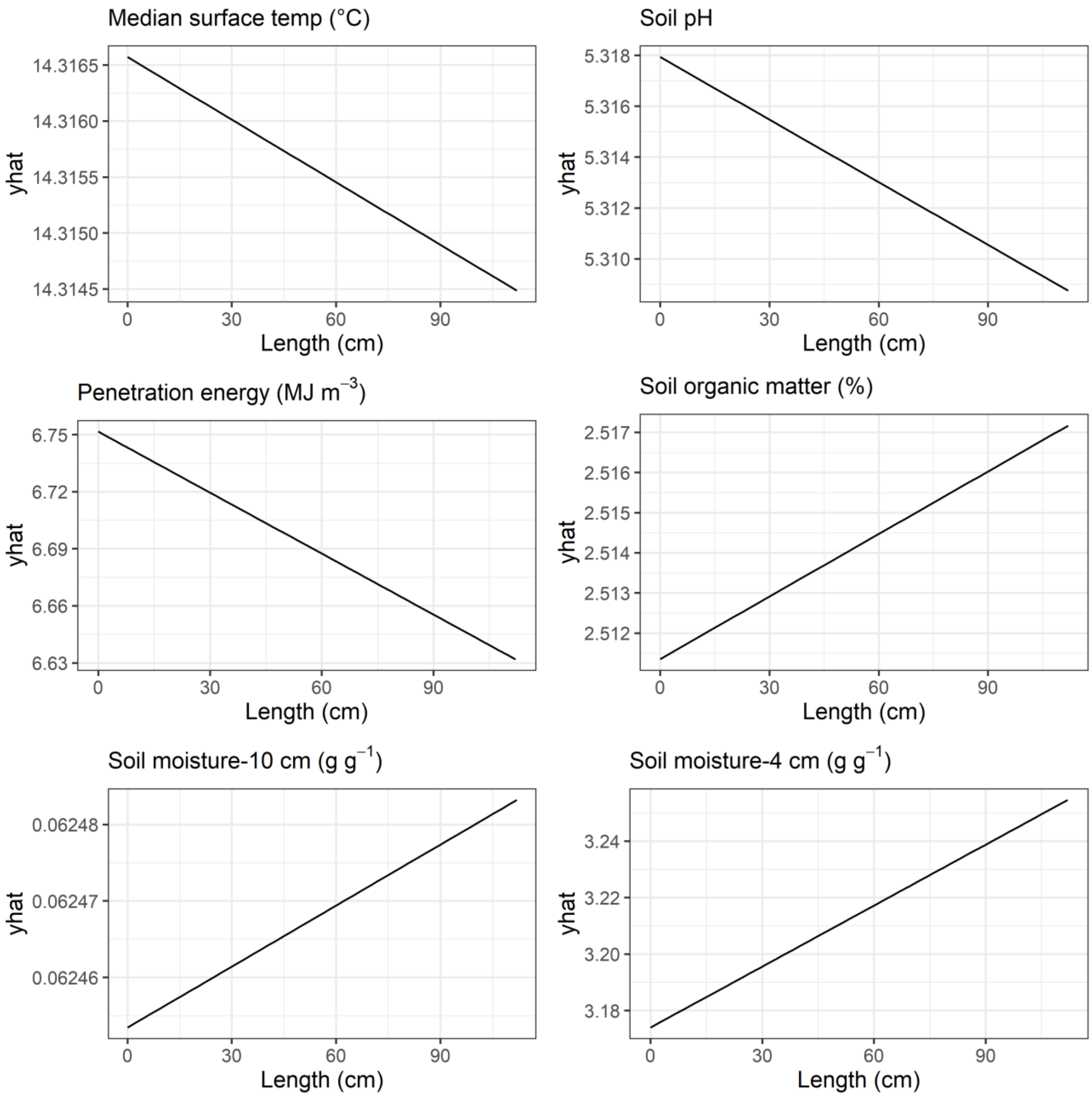

Figure S10: Partial dependence plots from six separate Generalized Linear Models (GLMs) for the response of environmental variables (median surface temperature (°C), soil pH, soil penetration energy (MJ $m^{-3}$), soil organic matter (%), soil moisture at 10 cm depth (g g-1), and soil moisture at 4 cm depth (g g-1)) to plant length (cm), taxon (n=17), plot, and the interaction between plant length and taxon. Plant length and location data were from the 2015 annual census, while environmental data for each plant location were extracted from kriged spatial data (10 cm resolution) from Blonder *et al.* (2018). Y-axis values are the average predicted values of the environmental variable based on plant length with other model predictors static, separated by taxon.

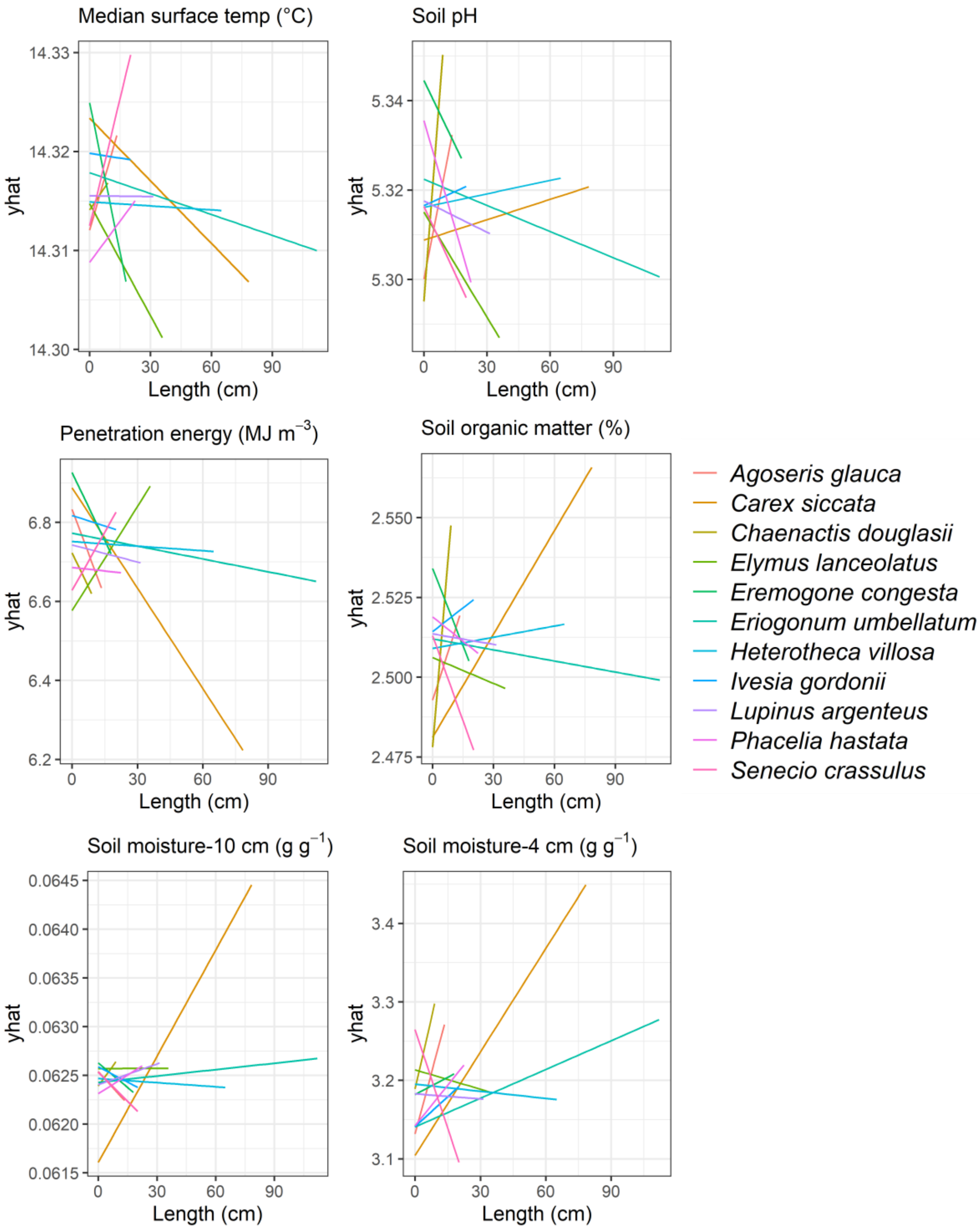

Figure S11: Parameter estimates for the random effects (Taxon and Plot) in the **survival** GLMM using **aboveground** estimates of overlap. A unique numeric plot identifier (1-50) was used to name plots.

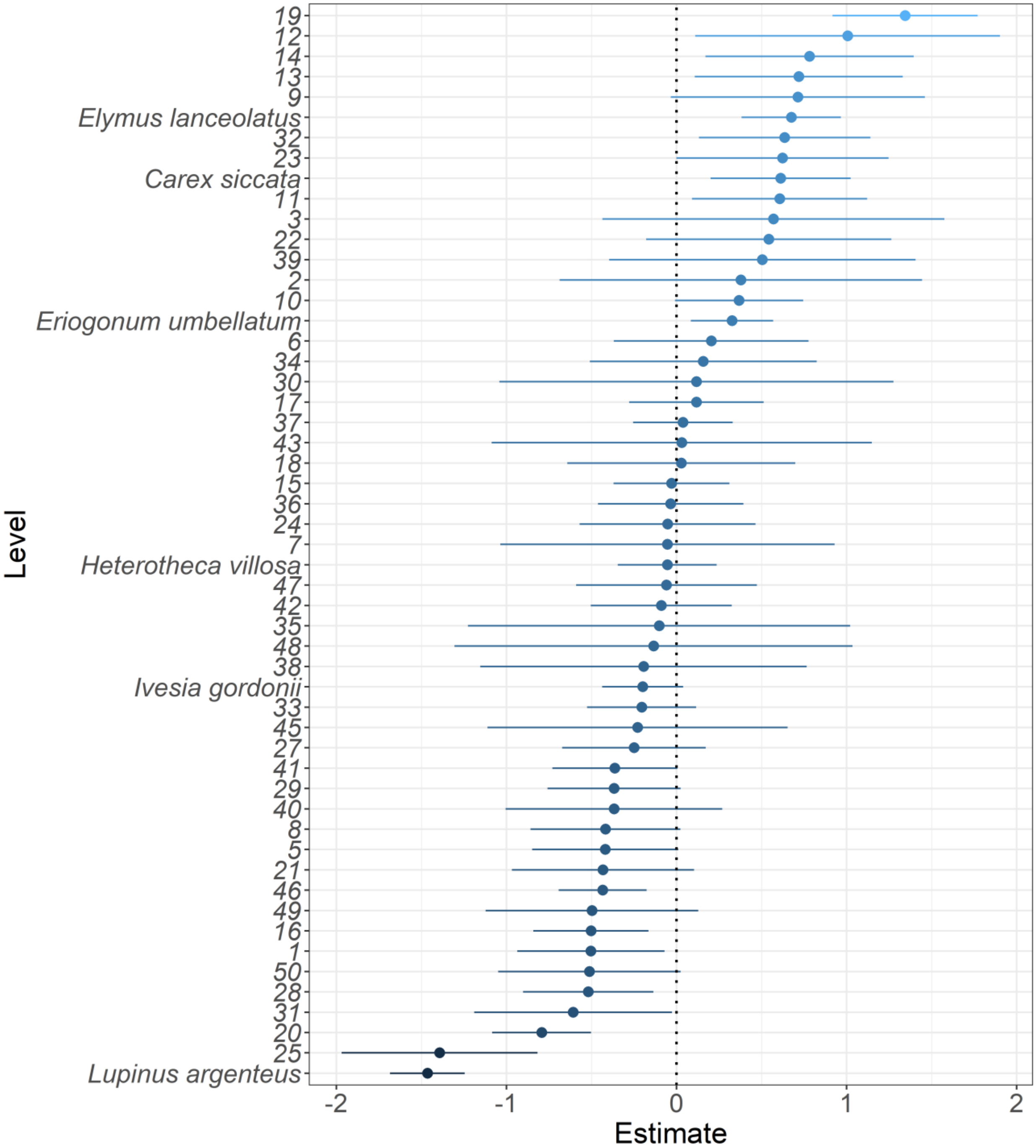

Figure S12: Parameter estimates for the random effects (Taxon and Plot) in the **survival** GLMM using **belowground** estimates of overlap. A unique numeric plot identifier (1-50) was used to name plots.

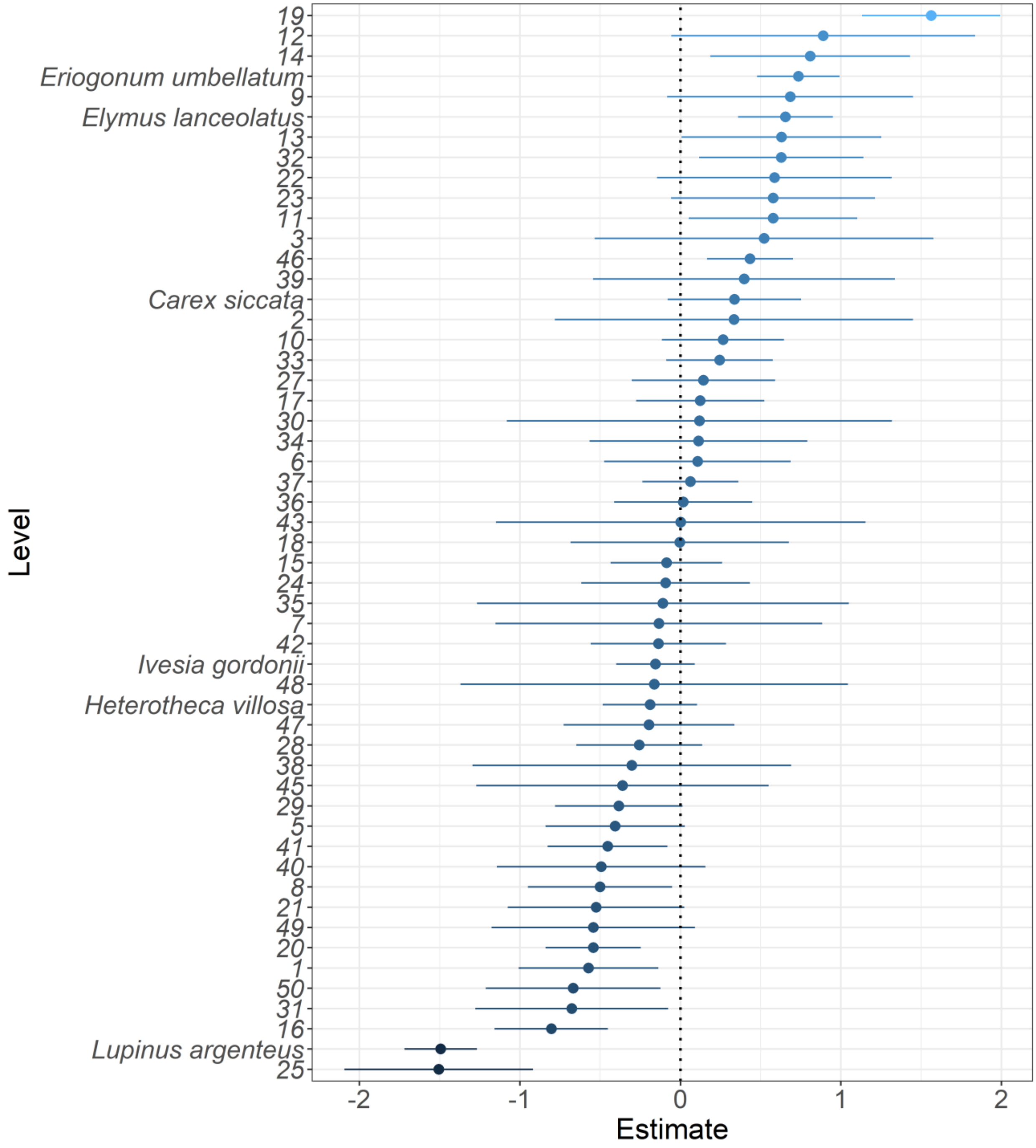

Figure S13: Parameter estimates for the random effects (Taxon and Plot) in the **growth** GLMM using **aboveground** estimates of overlap. A unique numeric plot identifier (1-50) was used to name plots.

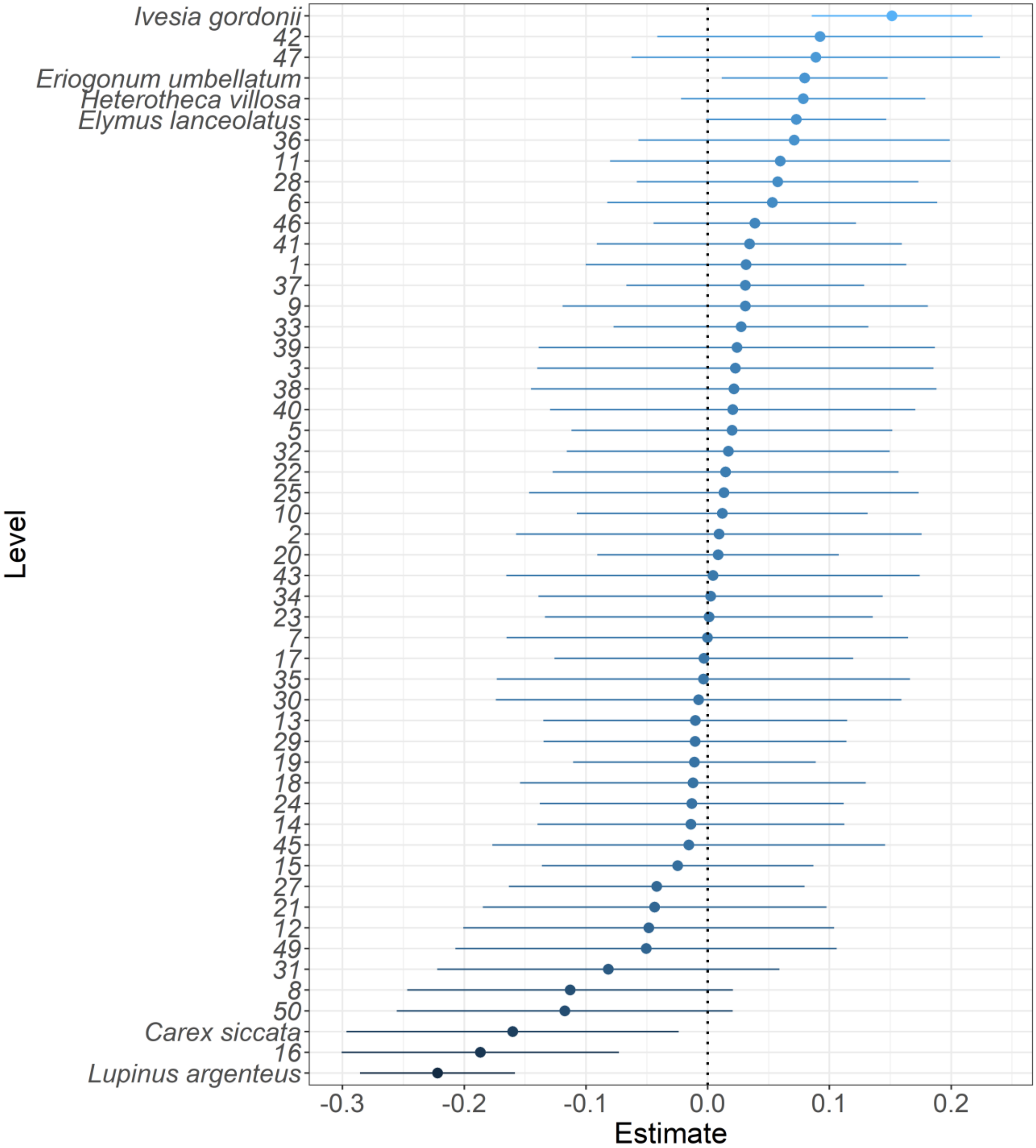

Figure S14: Parameter estimates for the random effects (Taxon and Plot) in the **growth** GLMM using **belowground** estimates of overlap. A unique numeric plot identifier (1-50) was used to name plots.

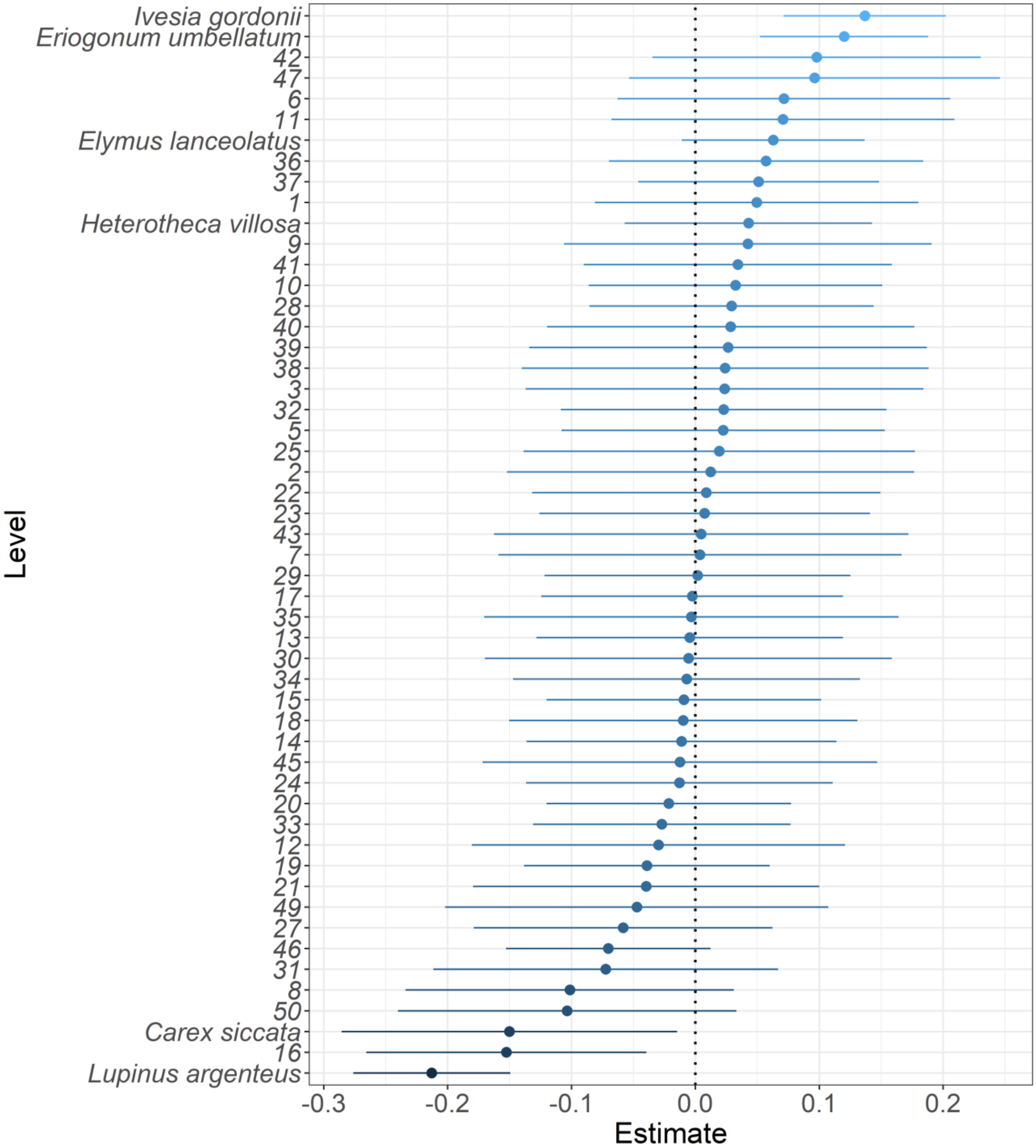

Figure S15: Parameter estimates for the random effects (Taxon and Plot) in the GLMM of **flowering probability** using **aboveground** estimates of overlap. A unique numeric plot identifier (1-50) was used to name plots.

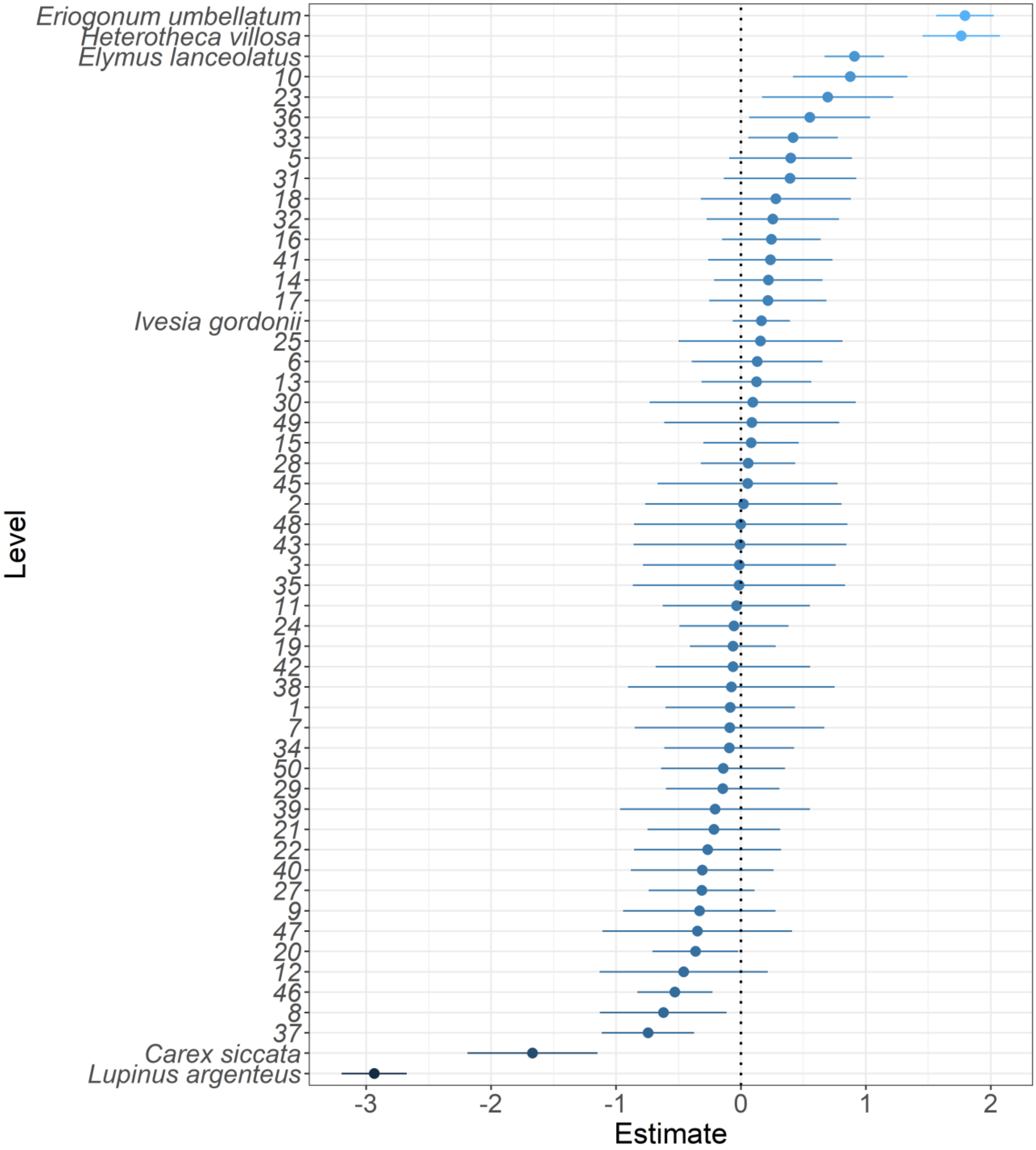

Figure S16: Parameter estimates for the random effects (Taxon and Plot) in the GLMM of **flowering probability** using **belowground** estimates of overlap. A unique numeric plot identifier (1-50) was used to name plots.

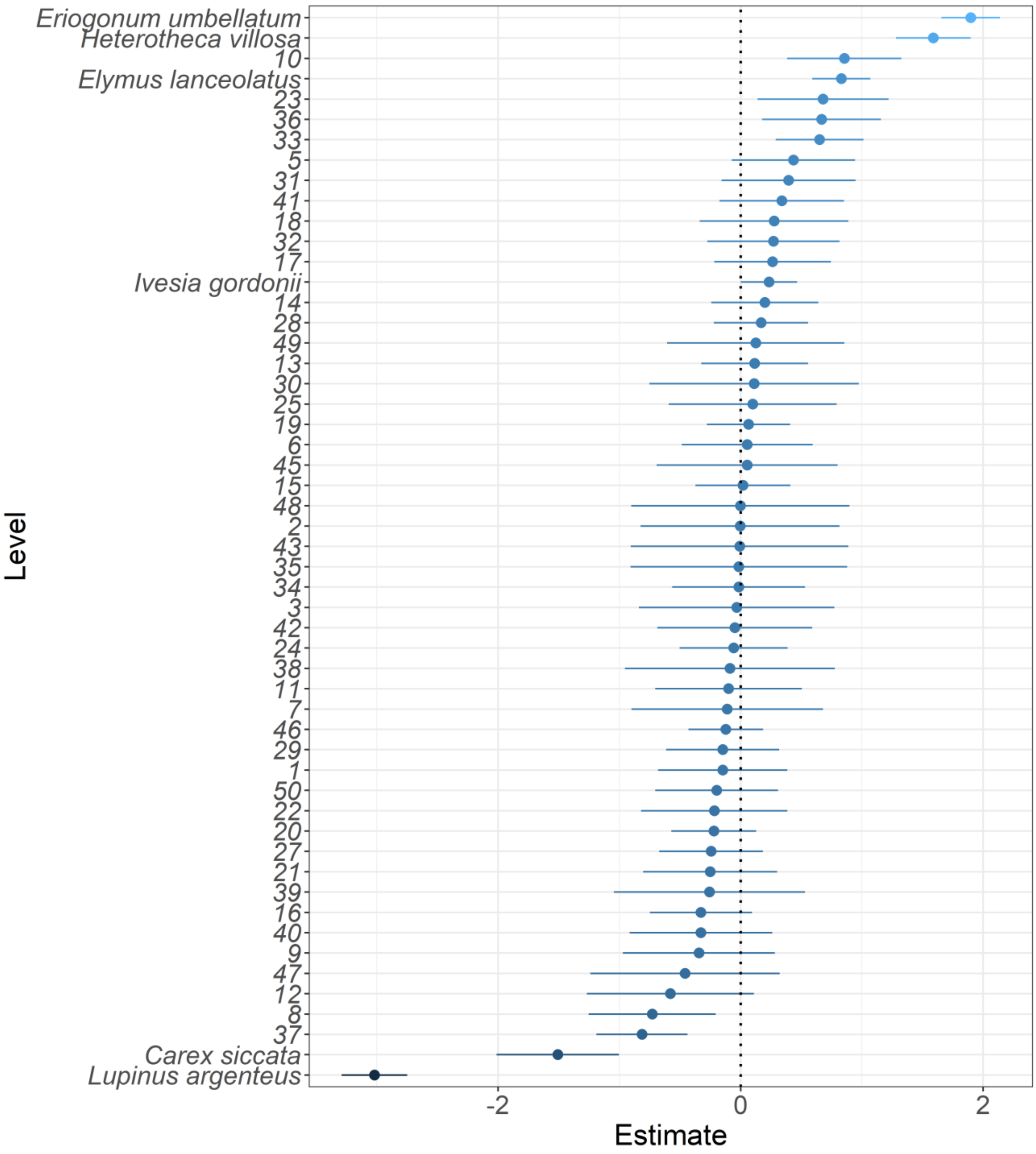

Figure S17: Parameter estimates for the random effects (Taxon and Plot) in the GLMM of log **number of inflorescences** using **aboveground** estimates of overlap. A unique numeric plot identifier (1-50) was used to name plots.

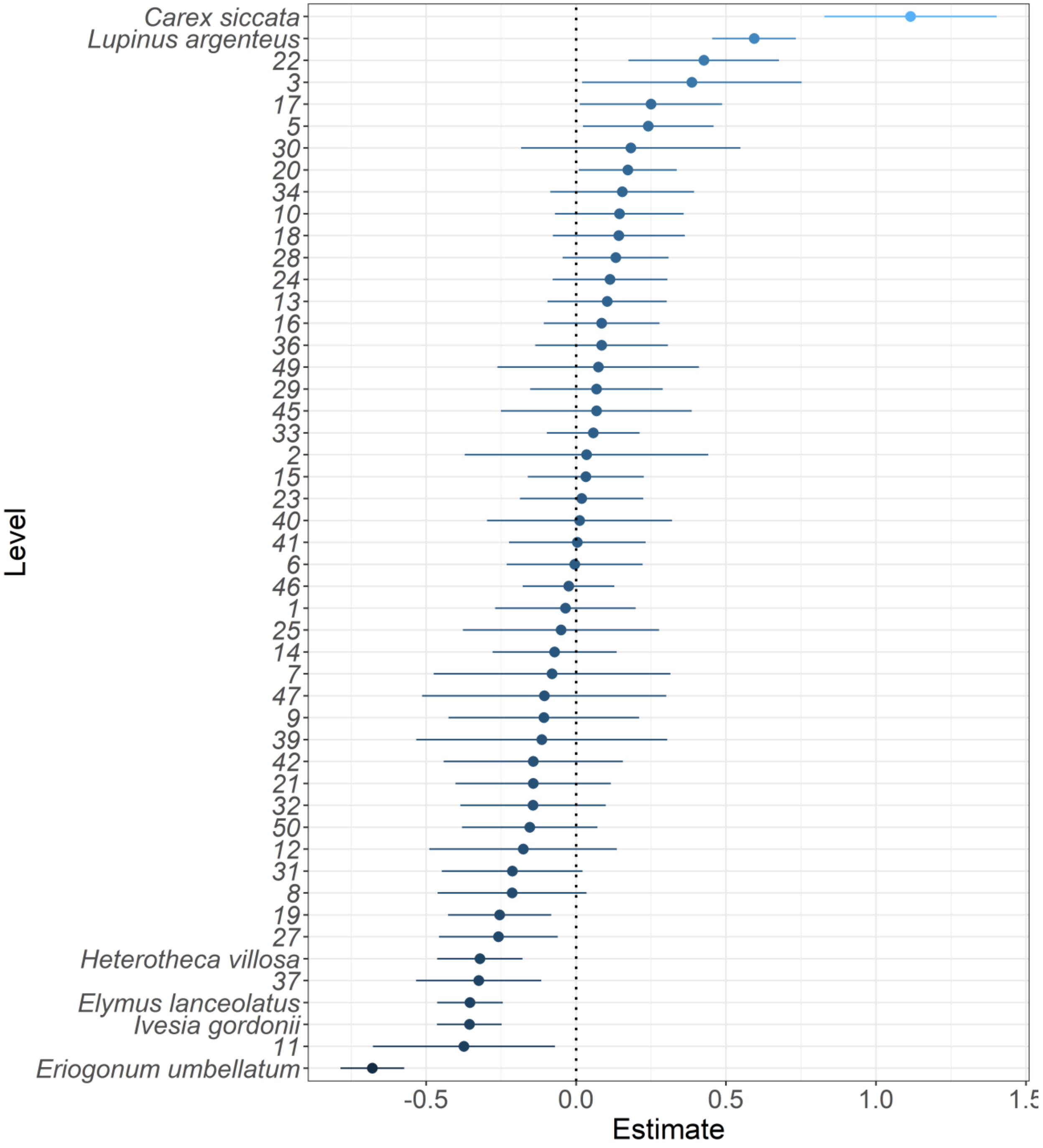

Figure S18: Parameter estimates for the random effects (Taxon and Plot) in the GLMM of log **number of inflorescences** using **belowground** estimates of overlap. A unique numeric plot identifier (1-50) was used to name plots.

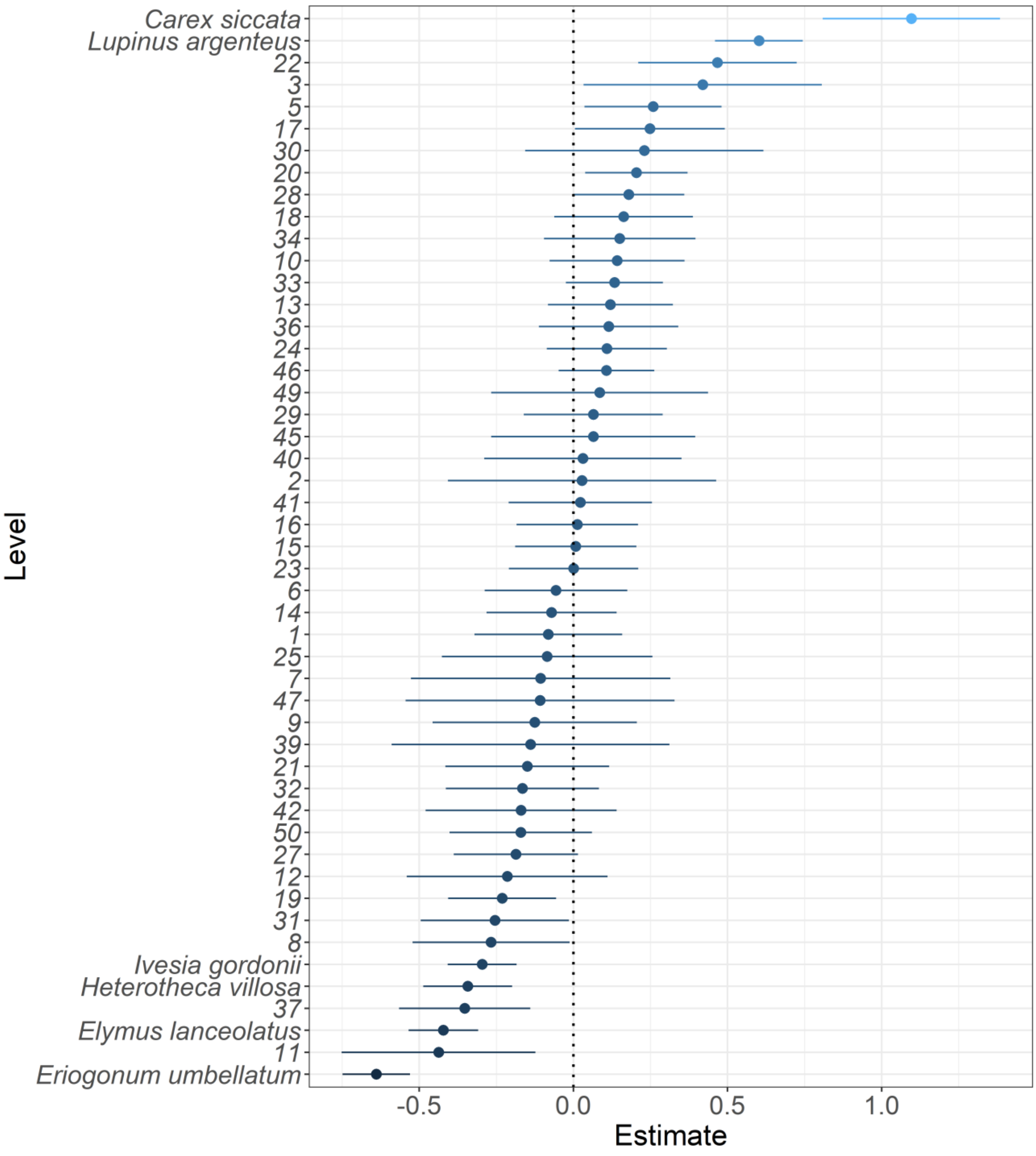

124 Table S1: Coefficient estimates with 95% confidence intervals for the **survival** GLMM using
125 **aboveground** estimates of percent overlap. Confidence intervals that exclude 1 are significant.

| | Survival | | |
|---|---|---|---|
| *Predictors* | *Odds Ratios* | *CI* | *p* |
| (Intercept) | 4.76 | 2.52 – 8.97 | **<0.001** |
| Precip | 1.47 | 1.37 – 1.58 | **<0.001** |
| Size | 1.98 | 1.82 – 2.15 | **<0.001** |
| Inter percent overlap | 0.91 | 0.83 – 1.00 | **0.044** |
| Intra percent overlap | 0.97 | 0.90 – 1.04 | 0.391 |
| Precip * Size | 0.88 | 0.81 – 0.95 | **0.001** |
| Precip * Inter percent overlap | 0.96 | 0.89 – 1.04 | 0.350 |
| Size * Inter percent overlap | 1.10 | 1.00 – 1.21 | 0.054 |
| Precip * Intra percent overlap | 0.96 | 0.89 – 1.03 | 0.289 |
| Size * Intra percent overlap | 1.26 | 1.15 – 1.39 | **<0.001** |
| (Precip * Size) * Inter percent overlap | 1.00 | 0.91 – 1.10 | 0.965 |
| (Precip * Size) * Intra percent overlap | 0.99 | 0.90 – 1.08 | 0.782 |
| **Random Effects** | | | |
| $\sigma^2$ | 3.29 | | |
| $\tau_{00\ Plot}$ | 0.39 | | |
| $\tau_{00\ Taxon}$ | 0.55 | | |
| ICC | 0.22 | | |
| $N_{Taxon}$ | 6 | | |
| $N_{Plot}$ | 47 | | |
| Observations | 5852 | | |
| Marginal $R^2$ / Conditional $R^2$ | 0.143 / 0.333 | | |

Table S2: Coefficient estimates with 95% confidence intervals for the **survival** GLMM using **belowground** estimates of percent overlap. Confidence intervals that exclude 1 are significant.

| | Survival | | |
|---|---|---|---|
| *Predictors* | *Odds Ratios* | *CI* | *p* |
| (Intercept) | 4.11 | 2.13 – 7.95 | **<0.001** |
| Precip | 1.52 | 1.41 – 1.64 | **<0.001** |
| Size | 1.96 | 1.79 – 2.13 | **<0.001** |
| Inter percent overlap | 0.72 | 0.65 – 0.81 | **<0.001** |
| Intra percent overlap | 0.90 | 0.81 – 1.00 | 0.052 |
| Precip * Size | 0.87 | 0.80 – 0.94 | **0.001** |
| Precip * Inter percent overlap | 1.10 | 1.02 – 1.18 | **0.010** |
| Size * Inter percent overlap | 0.99 | 0.92 – 1.07 | 0.843 |
| Precip * Intra percent overlap | 1.02 | 0.92 – 1.12 | 0.738 |
| Size * Intra percent overlap | 1.64 | 1.47 – 1.82 | **<0.001** |
| (Precip * Size) * Inter percent overlap | 1.08 | 1.00 – 1.18 | 0.060 |
| (Precip * Size) * Intra percent overlap | 1.04 | 0.94 – 1.15 | 0.453 |
| **Random Effects** | | | |
| $\sigma^2$ | 3.29 | | |
| $\tau_{00\ Plot}$ | 0.42 | | |
| $\tau_{00\ Taxon}$ | 0.59 | | |
| ICC | 0.23 | | |
| $N_{Taxon}$ | 6 | | |
| $N_{Plot}$ | 47 | | |
| Observations | 5852 | | |
| Marginal $R^2$ / Conditional $R^2$ | 0.218 / 0.401 | | |

Table S3: Coefficient estimates with 95% confidence intervals for the **growth** GLMM using **aboveground** estimates of percent overlap. Confidence intervals that exclude 0 are significant.

| | **Growth** | | |
|---|---|---|---|
| *Predictors* | *Estimates* | *CI* | *p* |
| (Intercept) | 0.08 | -0.06 – 0.21 | 0.259 |
| Precip | -0.01 | -0.04 – 0.02 | 0.541 |
| Size | 0.25 | 0.22 – 0.28 | **<0.001** |
| Inter percent overlap | 0.01 | -0.02 – 0.05 | 0.489 |
| Intra percent overlap | 0.01 | -0.02 – 0.05 | 0.420 |
| Precip * Size | 0.02 | -0.01 – 0.05 | 0.262 |
| Precip * Inter percent overlap | 0.07 | 0.04 – 0.10 | **<0.001** |
| Size * Inter percent overlap | -0.05 | -0.08 – -0.01 | **0.007** |
| Precip * Intra percent overlap | -0.00 | -0.03 – 0.02 | 0.744 |
| Size * Intra percent overlap | 0.04 | 0.01 – 0.08 | **0.024** |
| (Precip * Size) * Inter percent overlap | -0.00 | -0.04 – 0.03 | 0.822 |
| (Precip * Size) * Intra percent overlap | -0.02 | -0.05 – 0.01 | 0.136 |
| **Random Effects** | | | |
| $\sigma^2$ | 0.76 | | |
| $\tau_{00\ Plot}$ | 0.01 | | |
| $\tau_{00\ Taxon}$ | 0.03 | | |
| ICC | 0.04 | | |
| $N_{Taxon}$ | 6 | | |
| $N_{Plot}$ | 46 | | |
| Observations | 3737 | | |
| Marginal $R^2$ / Conditional $R^2$ | 0.074 / 0.112 | | |

Table S4: Coefficient estimates with 95% confidence intervals for the **growth** GLMM using **belowground** estimates of percent overlap. Confidence intervals that exclude 0 are significant.

| | **Growth** | | |
|---|---|---|---|
| *Predictors* | *Estimates* | *CI* | *p* |
| (Intercept) | 0.09 | -0.04 – 0.22 | 0.192 |
| Precip | -0.01 | -0.03 – 0.02 | 0.708 |
| Size | 0.24 | 0.21 – 0.27 | **<0.001** |
| Inter percent overlap | 0.09 | 0.05 – 0.13 | **<0.001** |
| Intra percent overlap | -0.01 | -0.05 – 0.03 | 0.667 |
| Precip * Size | 0.02 | -0.01 – 0.05 | 0.219 |
| Precip * Inter percent overlap | 0.04 | 0.01 – 0.07 | **0.011** |
| Size * Inter percent overlap | -0.02 | -0.05 – 0.00 | 0.089 |
| Precip * Intra percent overlap | 0.00 | -0.03 – 0.03 | 0.980 |
| Size * Intra percent overlap | -0.02 | -0.05 – 0.01 | 0.170 |
| (Precip * Size) * Inter percent overlap | -0.02 | -0.04 – 0.01 | 0.137 |
| (Precip * Size) * Intra percent overlap | -0.00 | -0.03 – 0.03 | 0.912 |
| **Random Effects** | | | |
| $\sigma^2$ | 0.76 | | |
| $\tau_{00\ Plot}$ | 0.01 | | |
| $\tau_{00\ Taxon}$ | 0.02 | | |
| ICC | 0.04 | | |
| $N_{Taxon}$ | 6 | | |
| $N_{Plot}$ | 46 | | |
| Observations | 3737 | | |
| Marginal $R^2$ / Conditional $R^2$ | 0.078 / 0.113 | | |

132 Table S5: Coefficient estimates with 95% confidence intervals for the **flowering probability** GLMM
133 using **aboveground** estimates of percent overlap. Confidence intervals that exclude 1 are significant.

| | **Flowering probability** | | |
|---|---|---|---|
| *Predictors* | *Odds Ratios* | *CI* | *p* |
| (Intercept) | 0.71 | 0.17 – 2.99 | 0.641 |
| Precip | 1.15 | 1.05 – 1.25 | **0.002** |
| Size | 10.15 | 8.79 – 11.71 | **<0.001** |
| Inter percent overlap | 0.84 | 0.75 – 0.94 | **0.002** |
| Intra percent overlap | 1.01 | 0.92 – 1.10 | 0.832 |
| Precip * Size | 1.12 | 1.01 – 1.26 | **0.040** |
| Precip * Inter percent overlap | 0.92 | 0.84 – 1.01 | 0.068 |
| Size * Inter percent overlap | 0.97 | 0.86 – 1.09 | 0.625 |
| Precip * Intra percent overlap | 1.04 | 0.96 – 1.13 | 0.336 |
| Size * Intra percent overlap | 0.96 | 0.86 – 1.06 | 0.412 |
| (Precip * Size) * Inter percent overlap | 0.96 | 0.85 – 1.07 | 0.446 |
| (Precip * Size) * Intra percent overlap | 1.05 | 0.95 – 1.17 | 0.301 |
| **Random Effects** | | | |
| $\sigma^2$ | 3.29 | | |
| $\tau_{00\ Plot}$ | 0.19 | | |
| $\tau_{00\ Taxon}$ | 3.14 | | |
| ICC | 0.50 | | |
| $N_{Plot}$ | 47 | | |
| $N_{Taxon}$ | 6 | | |
| Observations | 5076 | | |
| Marginal $R^2$ / Conditional $R^2$ | 0.461 / 0.732 | | |

Table S6: Coefficient estimates with 95% confidence intervals for the **flowering probability** GLMM using **belowground** estimates of percent overlap. Confidence intervals that exclude 1 are significant.

| | **Flowering probability** | | |
|---|---|---|---|
| *Predictors* | *Odds Ratios* | *CI* | *p* |
| (Intercept) | 0.70 | 0.17 – 2.91 | 0.627 |
| Precip | 1.10 | 1.01 – 1.20 | **0.032** |
| Size | 10.82 | 9.32 – 12.57 | **<0.001** |
| Inter percent overlap | 0.78 | 0.68 – 0.89 | **<0.001** |
| Intra percent overlap | 1.18 | 1.03 – 1.35 | **0.016** |
| Precip * Size | 1.08 | 0.97 – 1.21 | 0.170 |
| Precip * Inter percent overlap | 1.06 | 0.98 – 1.16 | 0.146 |
| Size * Inter percent overlap | 0.80 | 0.72 – 0.88 | **<0.001** |
| Precip * Intra percent overlap | 0.96 | 0.86 – 1.07 | 0.450 |
| Size * Intra percent overlap | 1.54 | 1.34 – 1.78 | **<0.001** |
| (Precip * Size) * Inter percent overlap | 1.05 | 0.96 – 1.16 | 0.297 |
| (Precip * Size) * Intra percent overlap | 0.92 | 0.82 – 1.05 | 0.215 |
| **Random Effects** | | | |
| $\sigma^2$ | 3.29 | | |
| $\tau_{00\ Plot}$ | 0.21 | | |
| $\tau_{00\ Taxon}$ | 3.09 | | |
| ICC | 0.50 | | |
| $N_{Plot}$ | 47 | | |
| $N_{Taxon}$ | 6 | | |
| Observations | 5076 | | |
| Marginal $R^2$ / Conditional $R^2$ | 0.486 / 0.743 | | |

Table S7: Coefficient estimates with 95% confidence intervals for the log **number of inflorescences** GLMM using **aboveground** estimates of percent overlap. Confidence intervals that exclude 0 are significant.

| | **Log number of inflorescences** | | |
|---|---|---|---|
| *Predictors* | *Estimates* | *CI* | *p* |
| (Intercept) | -1.11 | -1.68 – -0.54 | **<0.001** |
| Precip | 0.02 | -0.03 – 0.06 | 0.442 |
| Size | 0.79 | 0.75 – 0.83 | **<0.001** |
| Inter percent overlap | -0.08 | -0.13 – -0.02 | **0.011** |
| Intra percent overlap | -0.04 | -0.10 – 0.01 | 0.106 |
| Precip * Size | -0.02 | -0.06 – 0.02 | 0.313 |
| Precip * Inter percent overlap | -0.06 | -0.11 – -0.01 | **0.015** |
| Size * Inter percent overlap | -0.05 | -0.10 – -0.00 | **0.035** |
| Precip * Intra percent overlap | 0.03 | -0.02 – 0.08 | 0.277 |
| Size * Intra percent overlap | -0.03 | -0.08 – 0.02 | 0.277 |
| (Precip * Size) * Inter percent overlap | 0.02 | -0.03 – 0.07 | 0.425 |
| (Precip * Size) * Intra percent overlap | 0.01 | -0.04 – 0.06 | 0.618 |
| **Random Effects** | | | |
| $\sigma^2$ | 0.72 | | |
| $\tau_{00\ Plot}$ | 0.05 | | |
| $\tau_{00\ Taxon}$ | 0.49 | | |
| ICC | 0.43 | | |
| $N_{Plot}$ | 43 | | |
| $N_{Taxon}$ | 6 | | |
| Observations | 1963 | | |
| Marginal $R^2$ / Conditional $R^2$ | 0.344 / 0.624 | | |

Table S8: Coefficient estimates with 95% confidence intervals for the log **number of inflorescences** GLMM using a **belowground** estimates of percent overlap. Confidence intervals that exclude 0 are significant.

| | **Log number of inflorescences** | | |
|---|---|---|---|
| *Predictors* | *Estimates* | *CI* | *p* |
| (Intercept) | -1.14 | -1.71 – -0.58 | <**0.001** |
| Precip | 0.01 | -0.03 – 0.06 | 0.547 |
| Size | 0.79 | 0.74 – 0.83 | <**0.001** |
| Inter percent overlap | -0.05 | -0.13 – 0.03 | 0.252 |
| Intra percent overlap | -0.11 | -0.17 – -0.04 | **0.001** |
| Precip * Size | -0.02 | -0.06 – 0.02 | 0.364 |
| Precip * Inter percent overlap | -0.06 | -0.11 – -0.00 | **0.041** |
| Size * Inter percent overlap | -0.08 | -0.13 – -0.04 | <**0.001** |
| Precip * Intra percent overlap | 0.06 | 0.00 – 0.11 | **0.036** |
| Size * Intra percent overlap | 0.07 | 0.02 – 0.12 | **0.005** |
| (Precip * Size) * Inter percent overlap | 0.05 | -0.00 – 0.09 | 0.057 |
| (Precip * Size) * Intra percent overlap | -0.00 | -0.05 – 0.05 | 0.989 |
| **Random Effects** | | | |
| $\sigma^2$ | 0.71 | | |
| $\tau_{00\ Plot}$ | 0.06 | | |
| $\tau_{00\ Taxon}$ | 0.48 | | |
| ICC | 0.43 | | |
| $N_{Plot}$ | 43 | | |
| $N_{Taxon}$ | 6 | | |
| Observations | 1963 | | |
| Marginal $R^2$ / Conditional $R^2$ | 0.348 / 0.628 | | |

142
